# Utilising affordable smartphones and open-source time-lapse photography for monitoring pollinators

**DOI:** 10.1101/2024.01.31.578173

**Authors:** Valentin Ștefan, Aspen Workman, Jared C. Cobain, Demetra Rakosy, Tiffany M. Knight

## Abstract

Monitoring plant-pollinator interactions is crucial for understanding factors that influence these relationships across space and time. While traditional methods in pollination ecology are time-consuming and resource-intensive, the growing availability of photographic technology, coupled with advancements in artificial intelligence classification, offers the potential for non-destructive and automated techniques. However, it is important that the photographs are of high enough quality to enable insects to be identified at lower taxonomic levels, preferably genus or species levels. This study assessed the feasibility of using smartphones to automatically capture images of insects visiting flowers and evaluated whether the captured images offered sufficient resolution for precise insect identification. Smartphones were positioned above target flowers from various plant species to capture time-lapse images of any flower visitor in urban green areas around Leipzig and Halle, Germany. We present the proportions of insect identifications achieved at different taxonomic levels, such as order, family, genus, and species, and discuss whether limitations stem from the automated approach (e.g., inability to observe distinguishing features in images despite high image quality) or low image quality. Practical recommendations are provided to address these challenges. Our results indicate that for bee families, nearly three quarters of all cases could be identified to genus level. Flies were more difficult, due to the small size of many individuals and the more challenging features needed for identification (e.g., in the wing veins). Overall, we suggest that smartphones are an effective tool when optimised by researchers. As technology continues to advance, smartphones are becoming increasingly accessible, affordable, and user-friendly, rendering them an appealing option for pollinator monitoring.

## INTRODUCTION

The interactions between plants and their animal pollinators provide critical ecosystem services, by sustaining the reproduction of the majority of our food and wild plant species (Ollerton et al. 2011). It is thus critical to monitor plant-pollinator interactions and to understand the factors that cause these interactions to change across space and time. As many species of pollinating insects cannot be identified on sight, most studies quantifying plant-pollinator interactions capture insects that are observed to visit flowers and identifying them later using microscopy (e.g., Motivans Švara et al. 2021; Rakosy et al. 2022) or DNA barcoding (Creedy et al. 2020). While these methods are accurate and can provide museum specimens that are valuable for a wide variety of research purposes (Rakosy et al. 2023), they are time consuming, costly to scale, and require expert knowledge in insect taxonomy or barcoding.

Further, these methods are destructive, requiring the killing of many insects in order to monitor biodiversity and interactions. With the growing availability of video and photographic devices and artificial intelligence identification approaches, there is an opportunity for this field of science to move more towards automated, non-destructive methods in pollinator research (Montero-Castaño et al. 2022).

Recent review studies show that there is great potential to automate the detection and identification of plant-pollinator interactions (Martineau et al. 2017; Barlow & O’Neill 2020; Pegoraro et al. 2020; Høye et al. 2021; Amarathunga et al. 2021). Automated methods require cameras that can capture images of insects on flowers in a field setting and machine learning approaches that can identify the insects in the images based on their distinguishing morphological features (e.g., Stark et al. 2023). However, not all pollinating insects can be identified to the species level from an image, as distinguishing features may not be captured (i.e. features on the underside of the animal, genitalia) and/or are only visible under a microscope (e.g., certain hairs or micropunctures). Thus, a goal for a camera system for pollination research is to collect images in the field that are high enough quality that taxonomists, and thus machines, can identify the insect to the lowest taxonomic grain that can be expected from an image.

There are several features that are important for camera systems monitoring pollinators. First, the camera must be able to capture sharp consecutive images of fast-moving insects of variable dimensions in constantly changing light conditions on complex/noisy flower backgrounds. Second, the camera must be able to withstand extreme temperatures and/or high levels of humidity. Third, the camera system must not run out of power during the field sampling (ideally, run for 8 hours or more). Fourth, the cameras must be able to capture an image without deterring pollinators from visiting (i.e., should not need to be placed too close to a flower in order to provide a focused image, and should not make noise). Fifth, the cameras should be affordable and easy to buy and repair. This is critical to allow researchers with limited funding to participate in automated pollinator monitoring. Sixth, the cameras should be easy to set up, maintain, and transport so that multiple sites can be monitored. Finally, the cameras should produce image data with minimal size and maximal quality to prevent storage problems.

The development of camera systems for biological monitoring is rapidly evolving and was recently reviewed by Pegoraro et al. (2020). Numerous technologies are under investigation for monitoring pollinators. For example, camera systems include off-the-shelf digital products such as Apple iPod nanos (Lortie et al. 2012), fixed-lens cameras (Steen 2017), camera traps (Mcelveen & Meyer 2020; Naqvi et al. 2022), time-lapse cameras (Edwards et al. 2015; Smith et al. 2021; Alison et al. 2022; Nagai et al. 2022), and surveillance cameras (Steen & Thorsdatter Orvedal Aase 2011; Mertens et al. 2021), as well as recent programmable microcomputers such as NVIDIA Jetson Nano (Bjerge et al. 2022), Raspberry Pi (Ratnayake et al. 2021, 2023; Droissart et al. 2021; Bjerge et al. 2023), a Luxonis microcomputer-camera capable of edge AI coupled with a Raspberry Pi (Sittinger et al. 2023) and near-infrared sensors, like those offered by FaunaPhotonics (Rydhmer et al. 2022).

In this study, we explored the use of smartphones and time-lapse photography to photograph diurnal insects visiting flowers. Smartphones offer several potential advantages, including: (1) affordability, as phones with decent cameras are inexpensive (e.g., 85-130 euros) and (2) can be supplemented with light and cost-efficient power banks and tripods. (3) Persons without programming skills can easily set up and use smartphones to capture pictures of insects visiting flowers. For example, time-lapse photography apps, such as ‘Open Camera’ (Harman 2023), are already available for Android OS smartphones. (4) There are no supply shortages as, due to high demand, smartphones are always available from big distributors (unlike microcomputers). (5) Additionally, smartphones are capable of monitoring various habitat variables through integrated sensors for sound, location, ambient light, atmospheric pressure, and temperature. This extends their utility beyond image capture, allowing for the measurement of the surrounding environment (Lahoz-Monfort & Magrath 2021). (6) Finally, the ubiquity of smartphones makes them an ideal tool for citizen science initiatives, broadening the scope and scale of ecological data collection.

Few studies have attempted to use smartphones for the purpose of monitoring pollinators. Donovan et al. (2021) investigated the potential of using smartphones and motion capture for wildlife monitoring, including the possibility of monitoring pollinators. However, the authors found their setup to be inefficient for capturing clear images of pollinators. Specifically, many of their images did not contain pollinators and appeared to have been triggered by wind and plant movement rather than by the insects themselves, making reliable identification impossible.

Ratnayake et al. (2021) employed a Samsung Galaxy S8 smartphone camera placed on a tripod, positioned at a height of 0.6 m above a patch of *Scaevola* flowers, to capture videos of honeybee visitors. They reported high success with this method, however their focus was exclusively on a quantitative assessment of honeybees visits to a single plant species. More studies are needed to assess whether smartphones can provide useful images that would allow for biodiversity monitoring and research on plant-pollinator interactions.

With this paper, we aim to assess the use of smartphones for automatically capturing images of insects visiting flowers. We first give a detailed description of the observational set up and our approach in image selection and taxonomic identification. Second, we quantify the grain of taxonomic identification that is possible from the images collected by having expert entomologists identify the insect to the lowest possible level. We report the proportion of individuals identified at different taxonomic levels such as order, family, genus, and species level. Next, we discuss if the limited identification at finer taxonomic levels was a limitation of the automated approach in general (i.e., observing distinguishing features is not possible from a high-quality image) or due to low image quality (i.e. low resolution, low depth of field). Finally, we communicate some of the lessons learned and troubleshooting that were involved with using smartphones for pollination research.

## MATERIAL AND METHODS

Our study was conducted in the urban green areas in and around Leipzig and Halle, Germany. These field sites were chosen because they were active areas of research on pollinator biodiversity monitoring and plant-pollinator interaction studies. The list of sites is provided in Appendix Tab. VIII. Within these field sites, individual plants in flower were chosen for image collection of their visiting pollinators. These plants were chosen somewhat haphazardly at each site, based on the plant species that seemed to be getting a reasonable frequency of visits, as our goal was to capture images of visiting insects. We targeted flowers that were open and on top and avoided plant species that had drooping flowers, as these would be more difficult to photograph with our smartphone set up. In total, we monitored 33 different plant species (Appendix Tab. II). Images were collected from July through September 2021.

### Smartphone setup

To monitor visiting pollinators, we affixed a smartphone above the flower or inflorescence (setup example in Fig. 1). Smartphones captured time-lapse images with a time step of approximately 1-1.5 seconds for one hour on each target flower. After each session, the smartphone was moved to another flower. Most images were captured between 9:00 AM and 2:00 PM, although a few were taken as early as 8:00 AM and as late as 5:00 PM (Appendix Fig. I). Observations were conducted on sunny or mostly sunny days.

**Figure 1.**
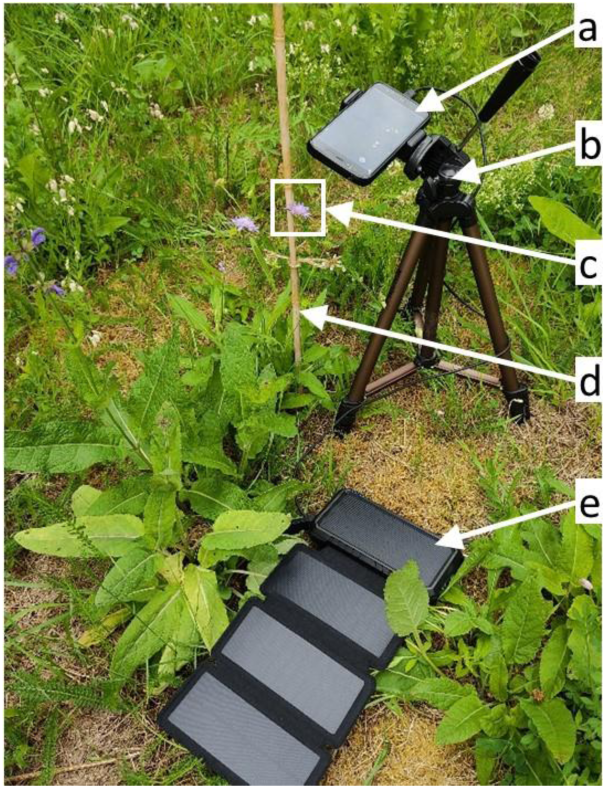
Example of the smartphone setup used for time-lapse photography, mounted on top of a target flower. a) smartphone, b) tripod, c) target flower, d) support stick used to stabilise the flower against wind movements, e) power bank.

The smartphones were securely mounted on tripods and powered through USB cables connected to power banks for a continuous flow of energy. We utilised power banks with an output of 5V and a range of 1-2.1mAh, resulting in a total cost of 130-200 EUR per unit (see gear example, phone models and costs in Appendix Tab. I). The gear setup was lightweight, weighing between 0.80 and 1.14 kg, with the power bank being the heaviest component, varying between 0.18 and 0.52 kg. We used the free Open Camera app (Harman 2023) for capturing the time-lapse images. The detailed protocols implemented for our field work can be found in Appendix Tab. VII.

To ensure consistent and high-quality recordings, we set a fixed focus at the start of each session on a target flower or portion of the flower/inflorescence which defined the region of interest. This was done by disabling the auto-focus feature in the Open Camera app, as it could result in images with a focus on the background rather than the target flower due to wind movements and insect activity. To minimise wind movements, we secured the target flowers to a wooden stick using yarn. The smartphones were positioned at a distance of 15-20 cm from the centre of the target flower to frame as much of a single flower as possible, which is important given the small size of most pollinators. We recorded each session for one hour, with the Open Camera app set to capture an image every second using time-lapse photography, and created a unique folder for each session with the plant’s name identified in the folder title. We used the Flora Incognita app (Mäder et al. 2021) to identify the focal plant species and this identification was verified by a botanist using the captured time-lapse images. At the end of the session, we manually stopped the recording. The Open Camera app allows the user to set the image resolutions, depending on the smartphone model. The majority of the images in our dataset were captured at a custom resolution of 1600 x 1200 pixels. The exposure was configured to automatic mode to adjust dynamically to changing lighting conditions.

### Insect annotation and identification

For each unique folder, we visually inspected each image to determine if it contained an insect. If an insect was present, we manually placed a tight bounding box around it and identified the insect to the taxonomic order. The free and open-source VGG Image Annotator (VIA) software (Dutta & Zisserman 2019) was utilised to view the images, draw bounding boxes, and enter the taxonomic order information. This software requires no installation, works across all common operating systems (Windows, Linux, or MacOS), and consists of a single HTML file that runs on most common web browsers (e.g., Google Chrome, Mozilla Firefox, etc.). To record annotation metadata for each bounding box, we created a custom JSON file template for VIA with a custom attribute table. A step-by-step annotation example and tutorial is provided on our GitHub repository (Ștefan 2023).

Next, we converted the JSON data files into spreadsheet files using R and Python scripts available on our GitHub repository. These spreadsheets contained the paths to images with insects, along with the pixel coordinates of the bounding boxes placed around the insects. Co-authors on this paper with entomological expertise visually inspected these images and identified the visitors in the Hymenoptera and Diptera orders to the lowest taxonomic level possible. Identification of insects within the order Hymenoptera was based on Westrich (2023). For Diptera we used Oosterbroek (2006), van Veen (2010) and Falk (2023). The identification of other visitors was limited to the order level only. To easily visualise each insect with its corresponding bounding box directly from the spreadsheets, we created the free and open source annotation tool ‘boxcel’ (Ștefan 2022).

## RESULTS AND DISCUSSION

We annotated 213 unique folders, which contained 33 different plant species (Appendix Tab. II) and 460,056 images. Each folder represented approximately one hour of time-lapse recording. We found 33,502 images (7.28%) that contained at least one insect, creating a total of 35,194 bounding boxes (the higher bounding box number was due to the presence of more than one insect in some images). Three annotators spent approximately 1,000 hours placing the bounding boxes and labelling the orders in the images. Taxonomists then spent additional time carrying out a detailed taxonomic identification, totalling an additional 720 hours. We estimate that it took 280 hours of fieldwork to collect the 213 annotated folders.

With the exception of one very small flower visitor for which only blurred images were captured, our smartphone-based approach enabled us to identify all insects to order level. In total we observed seven different orders of arthropods visiting our flowers (Tab. 1 & Fig. 2). Around 60% of the annotated bounding boxes contained a Hymenopteran, while 17% were Dipterans, with the two groups comprising more than three quarters of the annotated boxes. We rarely observed Lepidopterans (0.03%).

**Figure 2.**
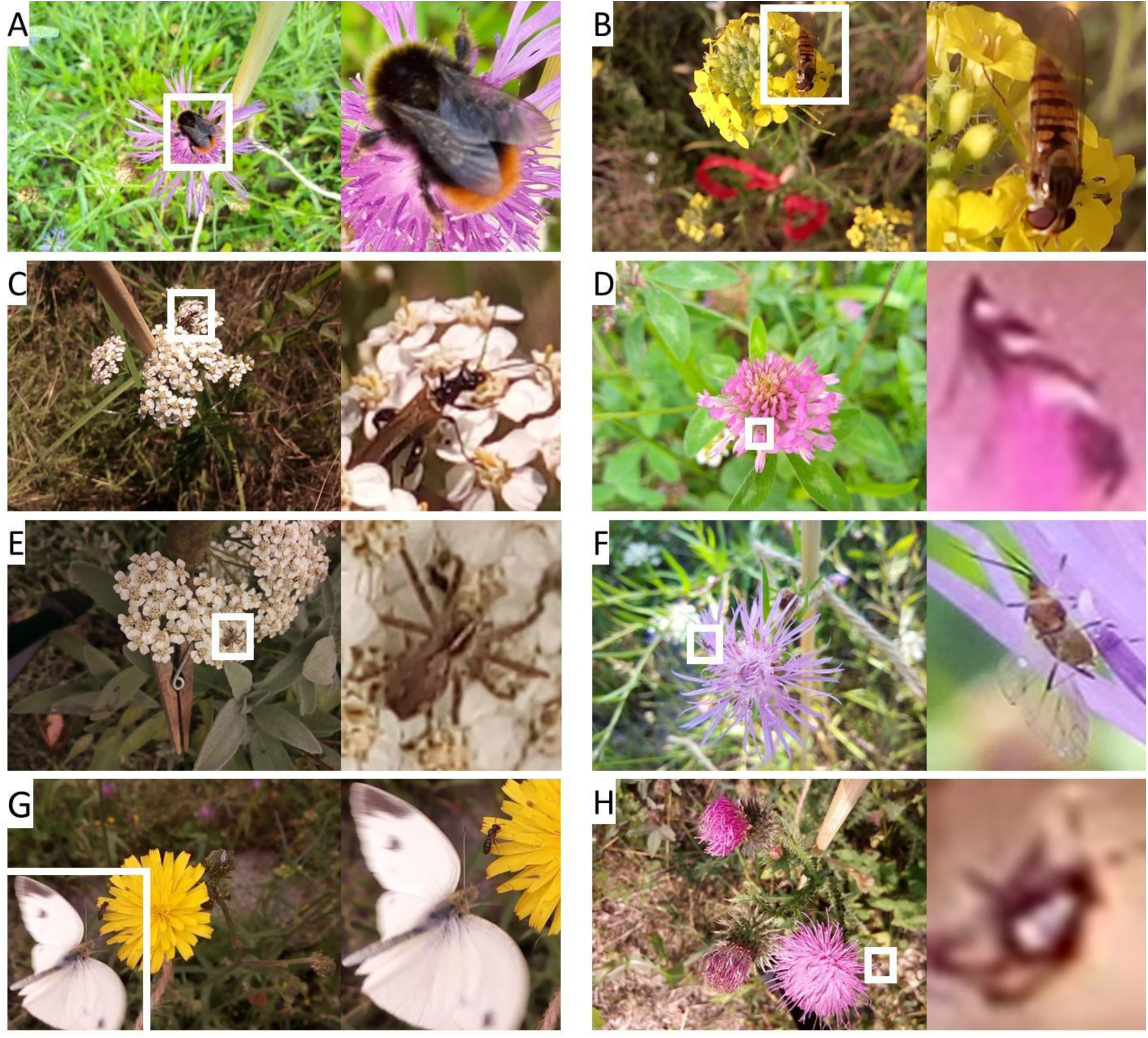
Examples of images with flower visitors from the seven observed orders with zoom on the respective bounding box for (A) Hymenoptera *- Bombus lapidarius* on flower of *Centaurea jacea*, (B) Diptera *- Episyrphus balteatus* on *Bunias orientalis*, (C) Coleoptera on *Achillea millefolium*, (D) Thysanoptera on *Trifolium pratense*, (E) Araneae on *Achillea millefolium*, (F) Hemiptera on *Centaurea jacea*, (G) Lepidoptera near *Picris hieracioides*, (H) No-Id, the order could not be identified.

**Table 1.** Number of annotated bounding boxes for each taxonomic order in the dataset. The 11 bounding boxes without taxonomic order labels (“No Id.”) correspond to 11 consecutive images of a very small flower visitor captured in blurred images.

| Order | No. box | Percent % | % cumul. sum |
| --- | --- | --- | --- |
| Hymenoptera | 20987 | 59.63 | 59.63 |
| Diptera | 5963 | 16.94 | 76.58 |
| Coleoptera | 3254 | 9.25 | 85.82 |
| Thysanoptera | 2965 | 8.42 | 94.25 |
| Araneae | 1158 | 3.29 | 97.54 |
| Hemiptera | 812 | 2.31 | 99.84 |
| Lepidoptera | 44 | 0.13 | 99.97 |
| No Id. | 11 | 0.03 | 100 |
| Total | 35194 | 100 |  |

### Identifying Hymenoptera to lower taxonomic levels

20,987 bounding boxes contained an individual in the order Hymenoptera (Appendix Tab. III & Fig. 3). Of these, 18,849 (89.8%) could be identified to one of ten families. There are a variety of reasons why some individuals could not be identified to the family level (Fig. 4). First, insects were outside the region of interest (ROI - Fig. 4A); in our case the ROI is typically a focal flower, an inflorescence, or a part of an inflorescence. Insects that were in the background rather than the ROI of the images were typically out of focus, obscured by surrounding vegetation, or appeared too small for further identification. For the purposes of many projects in pollination ecology, visitors on the focal flowers or inflorescence are the target of observations, and the ability to identify background insects that are outside this ROI would not be necessary. Second, identifying features of some insects that were inside the ROI were physically obscured from view in every image in which they occurred (Fig. 4B). For example, the insects were visiting the focal flower and the visible features of these insects clearly indicated they belonged in the order Hymenoptera (i.e. four wings, generally triangular-shaped head, striped abdomen, etc.), but the features necessary for identification to the family level were not visible. These insects were in clear focus, but were either partly cut out of the image due to movement of the flower after the camera was set up, obscured by the surrounding vegetation, or positioned at an angle such that only part of the insect was ever visible. Hereafter, these cases are simply referred to as “obscured”. Third, some insects within the ROI were too blurry to allow the discernment of the identifying features in every image in which they occurred (Fig. 4C). This occurred when the insect was too small to be seen in high resolution from the set depth of field, when the depth of field was too narrow and only focused on the surface of the flower but not the visiting insects, or when the insect was moving when the photo was taken. Hereafter, these cases are referred to as “too blurry.” Fourth, 1,655 bounding boxes were of a few individuals of tiny wasps that were only a handful of pixels in length (Fig. 4D). Because these tiny wasps remained on the same flower for several minutes at a time, we had hundreds of images of each individual. These tiny wasps were unlikely to be contributing to the pollination of the focal flowers and likely play a larger role in the ecosystem as parasitoids (Quicke 1997).

**Figure 3.**
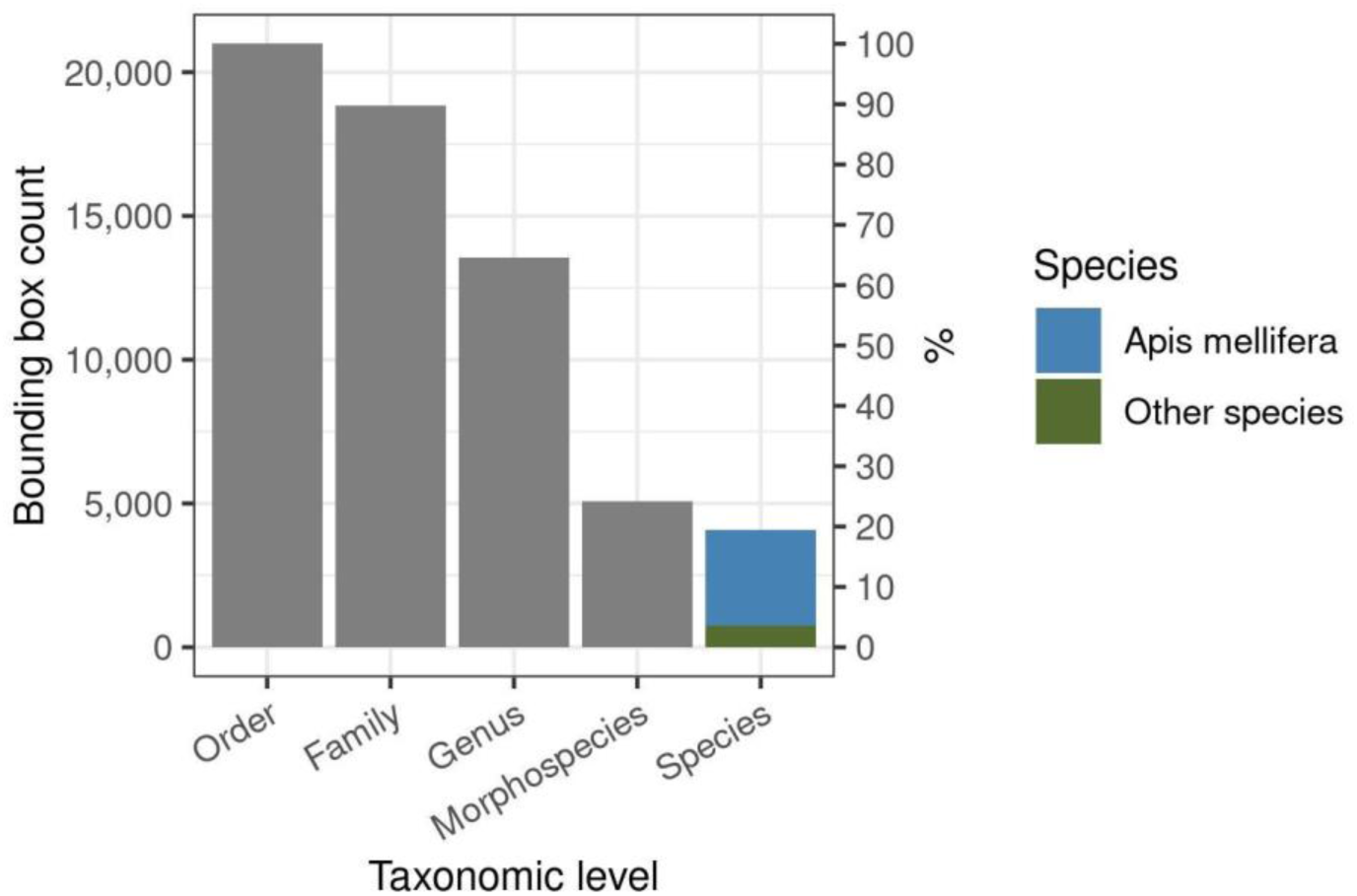
Bar plot illustrating the number and percent of bounding boxes that could be identified at each taxonomic level for the flower visitors in the order Hymenoptera. The primary y-axis (left) shows the count of boxes, while the secondary y-axis (right) displays the percentage of each taxonomic level relative to the total number of entries in the ‘Order’ level. This visualisation highlights the distribution and completeness of taxonomic information we could achieve within the Hymenoptera dataset. *Apis mellifera* comprised 15.8% of the total bounding boxes labelled as Hymenoptera (see also Appendix Tab. III).

**Figure 4.**
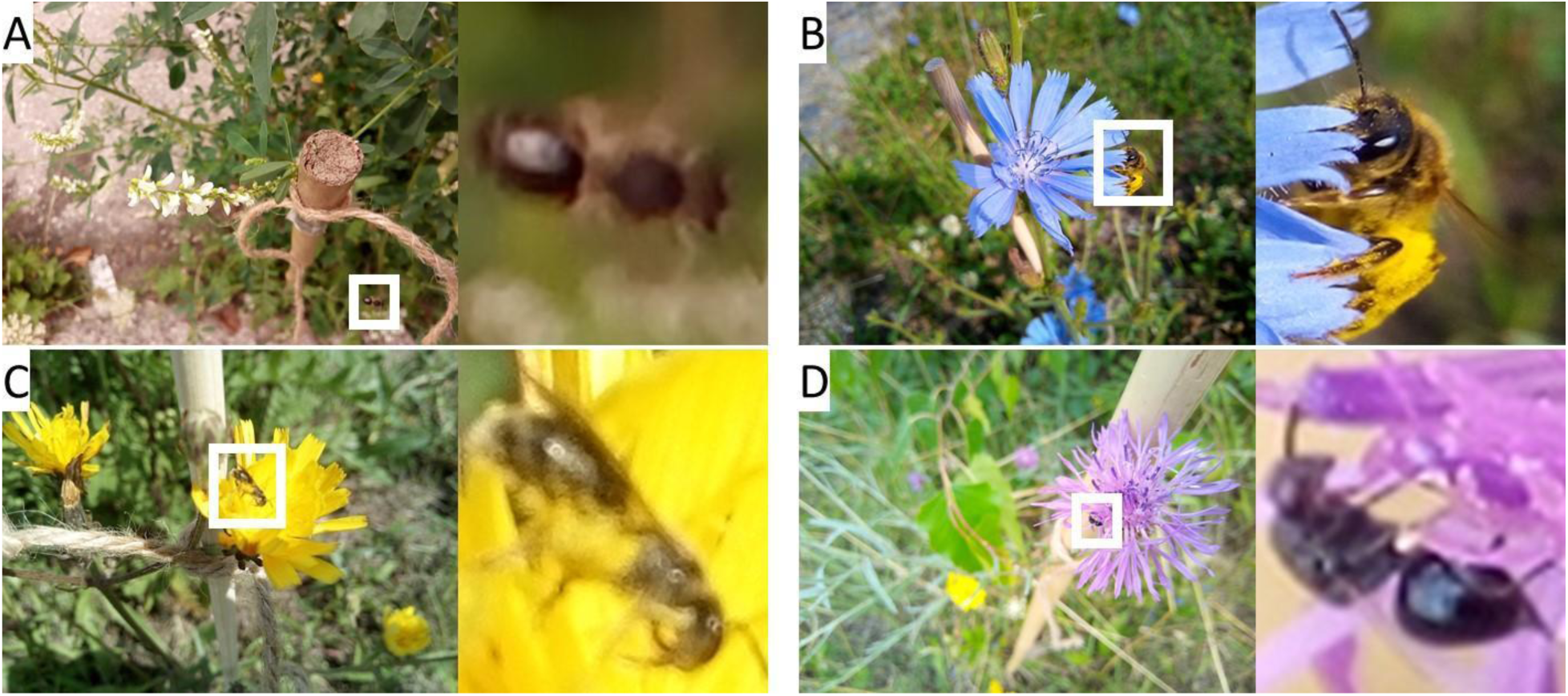
Examples of Hymenoptera insects that could not be identified to the family taxonomic level: (A) insect outside the region of interest (ROI), (B) insect within the ROI but all identifying features are physically obscured, (C) insect within the ROI but too blurry to identify, (D) tiny wasp.

The challenges of identifying very small insects are discussed further in the section Identifying Diptera to lower taxonomic levels. Of the 2,138 (10.2%) bounding boxes that could be identified to order Hymenoptera but not further identified to family, 397 (18.6%) were outside the region of interest, 11 (0.5%) were obscured, 76 (3.6%) were too blurry, and 1,655 (77.4%) were tiny wasps.

There were 608 bounding boxes that were identified to the family Andrenidae. Bees in this family within Germany are generally black and may have pale banding on the abdomen (Westrich 2023). Most species are the size of a honeybee (*Apis mellifera*) or smaller (Westrich 2023). Andrenid bees carry pollen in scopae on the hind tibiae and basal leg segments (Westrich 2023). Within the bounding boxes identified as Andrenidae, 253 bounding boxes could be further identified to the genus *Andrena* (Appendix Tab. III), due to the presence of facial foveae, patches of hairs between the compound eyes and the antennal socket (Fig. 5A), and the presence of three rather than two submarginal cells in the wings (Fig. 5B) (Westrich 2023). Another distinguishing trait is the hairs on the hind legs, used for pollen collection, although these were not visible in the images. Over 100 species of *Andrena* are present in Germany and determination to the species level is most reliant on dissection of the genitalia, rendering identification from images impossible. The remaining bounding boxes in the family Andrenidae could not be identified to genus level because a clear view of the bee’s face and wing venation are necessary to determine the genus. Of these bounding boxes that could not be identified to genus, one was outside the ROI, 28 were obscured, and 326 were too blurry.

**Figure 5.**
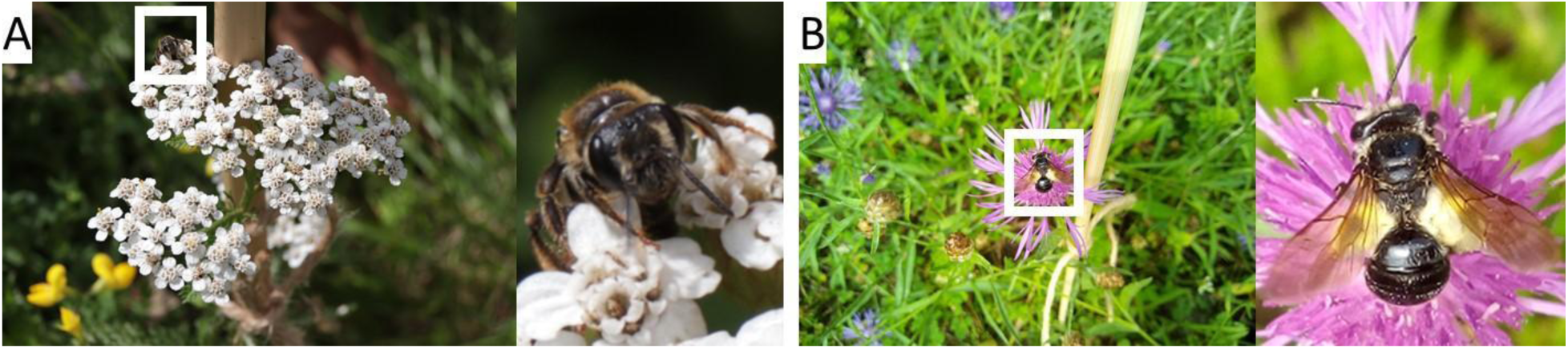
Examples of individuals from the *Andrena* genus: (A) showcasing distinctive facial foveae, (B) emphasising the presence of two submarginal cells, visible in the right wing’s venation.

There were 8,493 bounding boxes that were identified to the family Apidae, and 8,486 of these could be further identified to genus. The vast majority of these insects were *Apis mellifera* (3,307 individuals) or *Bombus* (5,178 individuals). A single bounding box was identified as an individual in the genus *Anthophora*, which was recognized by its green compound eyes, striped abdomen, and stocky body (Westrich 2023).

We captured seven bounding boxes containing a single bee from the Apidae family. While the images were too blurry to identify the bee down to the genus level, certain characteristics such as elongated marginal cells on the wings and a striped, stocky abdomen suggest it belongs to one of a few genera within the Apidae family (e.g., *Apis, Anthophora, Eucera, Tetralonia*). A clearer image of the wing venation to see the submarginal cells and a view of the scopae or corbiculae would have been necessary for further identification.

The Western honeybee, *Apis mellifera*, is easily identified by its characteristic golden-brown coloration, corbiculae (“pollen baskets”), striped abdomen, and banana-shaped marginal cell (Fig. 6A). These features are nearly always visible in the images.

**Figure 6.**
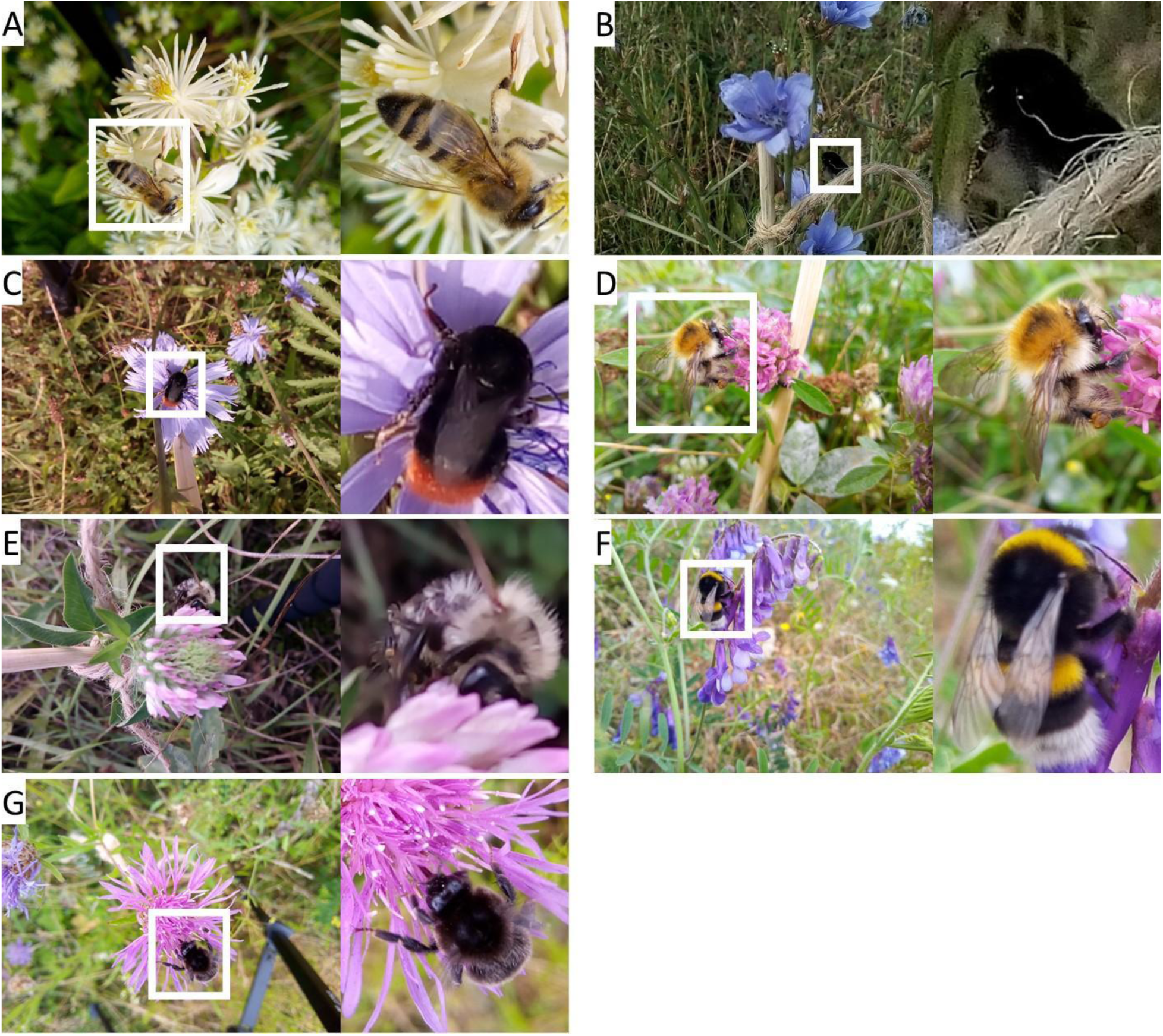
Examples of individuals from the Apidae family: (A) with all characteristic features of *Apis mellifera* - golden-brown coloration, corbiculae (“pollen baskets”), striped abdomen, and banana-shaped marginal cell, (B) example of *Bombus* “black” morphospecies group, (C) *Bombus* “red-tailed” morphospecies group, (D) *Bombus* “red and yellow” morphospecies group, (E) *Bombus* “striped” morphospecies group, (F) *Bombus* “white-tailed” morphospecies group, (G) *Bombus* individual which could not be assigned to a morphospecies group.

For the genus *Bombus*, species were grouped into five morphospecies groups which were defined by the shared coloration patterns of visually-similar species (Fig. 6B-F, Appendix Tab. IV). Identification to the species level within *Bombus* from our images alone is very challenging or even impossible due to shared morphology of cryptic species complexes, some of which can only be differentiated via genetic analysis. For most species with shared morphology, characteristics such as tongue length and features of the genitalia are used to differentiate between species, which are not visible in our images. The “black” morphospecies encompasses melanistic *B. hortorum* and *B. ruderatus*; these individuals are solid black with no other coloration. Seven bounding boxes contained individuals in the “black” morphospecies group. The “red-tailed” morphospecies is defined by the presence of red hairs on the terminal few abdominal segments and encompasses 12 taxa: *B. confusus confusus*, *B. cullumanus*, *B. lapidarius*, *B. monticola*, *B. pomorum*, *B. pratorum*, *B. pyrenaeus*, *B. ruderarius*, *B. rupestris*, *B. soroeensis proteus*, *B. sylvarum*, and *B. wurflenii*. It’s worth noting that *B. lapidarius* and *B. ruderarius* are more likely to be found in urban areas.

There were 3,676 bounding boxes that contained individuals in the “red-tailed” morphospecies group; seven of those bounding boxes were further identified as *B. lapidarius*, as they contained male bees with yellow hairs on the thorax but only black and red hairs on the abdomen. The “red and yellow” morphospecies group is defined by the presence of red hairs on the thorax and red and/or yellow hairs on the abdomen. This group encompasses four species: *B. humilis*, *B. muscorum*, *B. pascuorum*, and *B. schrencki*. There were 698 bounding boxes which contained individuals in the “red and yellow” morphospecies group. The “striped” morphospecies group is defined by one or two (generally quite bold) thoracic bands in yellow or white and variably-coloured abdominal banding. This group contains species which lack the strong “tail” marking of the other morphospecies groups and any red coloration on the thorax. This group encompasses 11 taxa: *B. armeniacus*, *B. campestris*, *B. distinguendus*, *B. fragrans*, *B. haematurus, B. mesomelas, B. mudicus, B. sichelii, B. subterraneus subterraneus, B. subterraneus latreillellus* (drones only) and*, B. veteranus*. There were ten bounding boxes which contained individuals in the “striped” morphospecies group. The “white-tailed” morphospecies group is defined by the presence of white hairs on the terminal few abdominal segments and represents the group which is the most unlikely to be differentiated by photos. This group encompasses 19 taxa: *B. argillaceus, B. barbutellus, B. bohemicus, B. confusus paradoxus, B. cryptarum, B. gerstaeckeri, B. hortorum, B. jonellus, B. lucorum, B. magnus, B. norvegicus, B. quadricolor, B. ruderatus, B. semenoviellus, B. soroeensis lectitatus, B. subterraneus latreillellus* (queens and workers only), *B. sylvestris, B. terrestris, and B. vestalis*. There were 672 bounding boxes which contained individuals in this “white-tailed” morphospecies group. Other species of Bombus which occur in Germany and are not covered by these morphospecies groups were categorized into three additional morphospecies groups (“black-tailed,” “red thorax,” and “primarily yellow”), but these groups were not captured in this annotated data set. An additional 115 bounding boxes contained individuals which could reliably be identified to the genus *Bombus*, but which were obscured such that the abdomen was not visible in any of the images (Fig. 6G). Without seeing the abdomen, it is not possible to place the individual into any of the morphospecies groups; these instances represent about 2% of the total occurrences of Bombus within this data set.

There were 404 bounding boxes that were identified to the family Colletidae, and all could be identified further to two genera that occur in Germany: *Colletes* and *Hylaeus*. The genus *Colletes* is most easily recognized by the S-shaped 2nd recurrent vein; however, this feature is rarely visible in the field images. Nevertheless, two bounding boxes were identified to the genus *Colletes* as they are clearly not *Hylaeus*. The genus *Hylaeus* is one of the most recognizable genera in the data set due to its distinctive morphology. Bees in this genus are fairly small, hairless, and black with white to yellow maculations, especially on the face and thorax. They also have two submarginal cells and the second is half the size of the first, which is unique to this genus when compared to other small, dark, hairless bees in Germany. There are nearly 40 species of *Hylaeus* that occur in Germany and differentiation between species depends on minute differences in the maculations that are not visible in the field images. There were 402 bounding boxes which were identified to the genus *Hylaeus*.

There were 390 bounding boxes that were identified to the family Cynipidae (gall wasps). The Cynipidae were easily recognized by their compact mesosoma, tall metasoma when viewed from the side, visible ovipositor, and small size. Lower-level identification of these individuals was not attempted, as we focus on pollinators rather than parasitoids (Whitfield 1998).

There were 2,426 bounding boxes that were identified to the family Formicidae (ants). The Formicidae were easily recognized by their lack of wings, their geniculate antennae, and the presence of a petiole between the mesosoma and metasoma. Lower-level identification of these individuals was not attempted due to our focus on pollinating insect groups. Ants are rarely pollinators in temperate grassland ecosystems, and more often disrupt pollination by damaging flowers or changing the foraging behaviour of pollinators (Wills & Landis 2018).

There were 6,153 bounding boxes that were identified to the family Halictidae. The primary trait which defines the Halictidae family is a strongly curved basal vein in the wing (Fig. 7A). This feature is not always visible in the field images, however, so it was often easier to look for features that define certain genera within Halictidae. The three most diverse genera within Halictidae in Germany are generally easy to identify to the genus level. For the genera *Halictus* and *Lasioglossum*, the characteristic trait is the presence of a “furrow” or division in the terminal abdominal segment of female individuals (Fig. 7B). This feature is generally very easy to see if the bee is positioned correctly. These two genera are further distinguished from each other by the positioning of the hairs on the abdominal segments: in *Halictus*, the hairs are positioned apically on the tergites, whereas in *Lasioglossum*, the hairs are positioned basally (Fig. 7C,D). Differentiating between these two positions is possible in higher quality images. A total of 2,959 bounding boxes were identified as individuals within the genus *Halictus* and 969 as individuals in the genus *Lasioglossum*. Further identification within these genera is complicated by the large number of species, but a few species are somewhat distinctive. *Halictus scabiosae* are distinguished by their large size and doubly-banded abdomen in females, but confusion with *H. sexcinctus* is possible, especially later in the season as the abdominal bands become worn. There were 287 bounding boxes that were identified as *H. scabiosae*.

**Figure 7.**
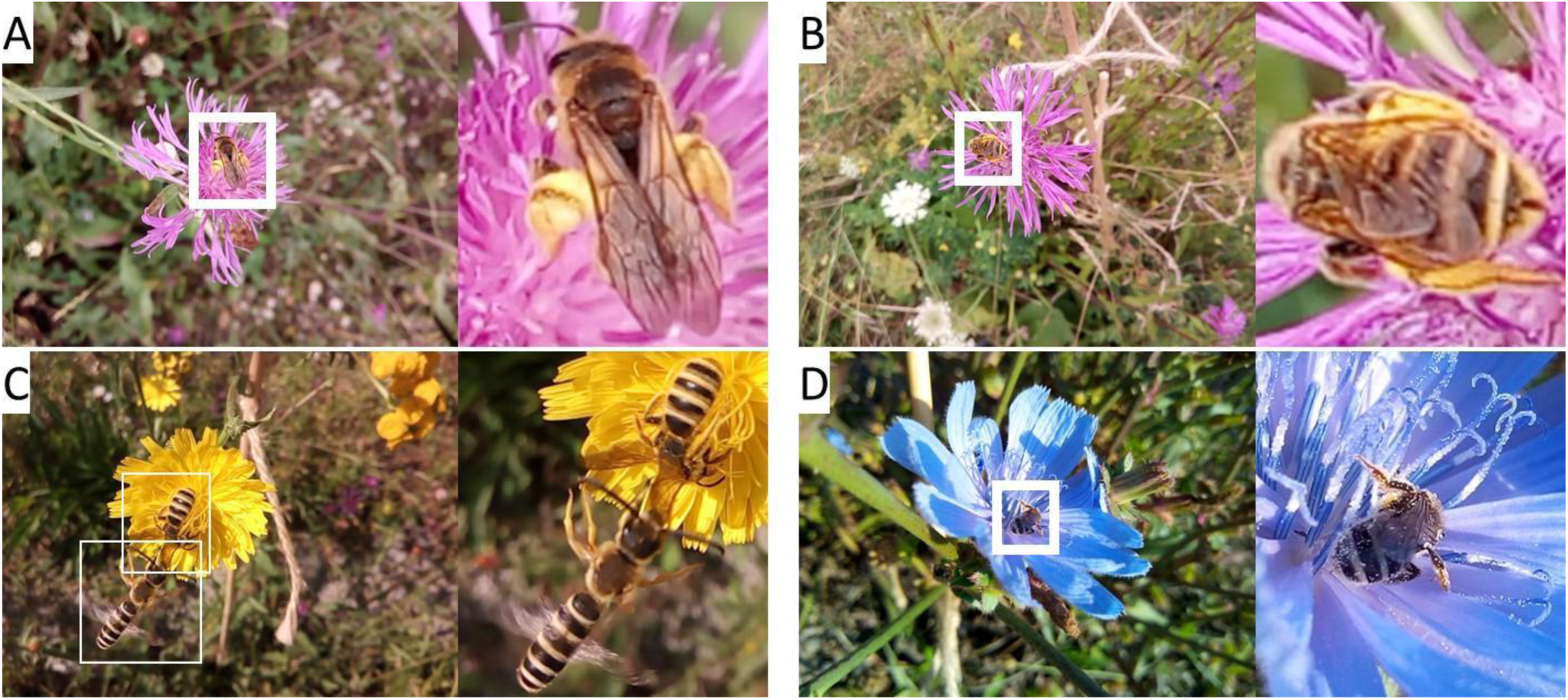
Examples of individuals from the Halictidae family: (A) with strongly curved basal wing vein characteristic of the family Halictidae, (B) with abdominal furrow, characteristic of females in the genera *Halictus* and *Lasioglossum*, (C) with apical banding of the tergites in *Halictus* male (lower individual) and female (upper individual), (D) with basal banding of the tergites in *Lasioglossum*.

Differentiation of the many small, golden-coloured *Halictus* can be challenging, but *H. subauratus* may be distinguished by their iridescent green compound eyes. There were 422 bounding boxes which were identified as *H. subauratus*. Males of certain *Lasioglossum* species can be distinguished by the red coloration of the abdomen, such as in *L. calceatum* and *L. interruptum*. There were 28 bounding boxes which were identified as *L. calceatum*. Bees within the genus *Sphecodes*, known as blood bees, belong to the third most diverse genus within the family Halictidae. They are distinguished by their red abdomen, hairless body and the lack of pollen-carrying scopa (all species are kleptoparasites which do not collect pollen). A total of 133 bounding boxes were identified as individuals within the genus *Sphecodes*. Differentiation between species within *Sphecodes* from images is nearly impossible as most species have the same black and red coloration and can vary in size within the same species.

There were 2,092 bounding boxes which were identified as individuals within the family of Halictidae, mostly due to the presence of the strongly curved basal vein or the presence of the abdominal furrow, but which could not be identified to genus. Of these boxes, 678 were outside the ROI, 142 were obscured, and 1,272 were too blurry to be identified to the genus level.

There were 187 bounding boxes which were identified to the family Megachilidae. This family is most easily distinguished by the ventral abdominal scopa in most genera. Two individuals (13 bounding boxes) were identified as *Anthidium manicatum*. This species is recognizable by its large size and broken yellow abdominal bands (Fig. 8A). Differentiating between two of the other more common genera in Germany, *Megachile* and *Osmia*, is based on the presence (*Osmia*) or absence (*Megachile*) of arolia between the claws, a trait that is not visible in our images.

**Figure 8.**
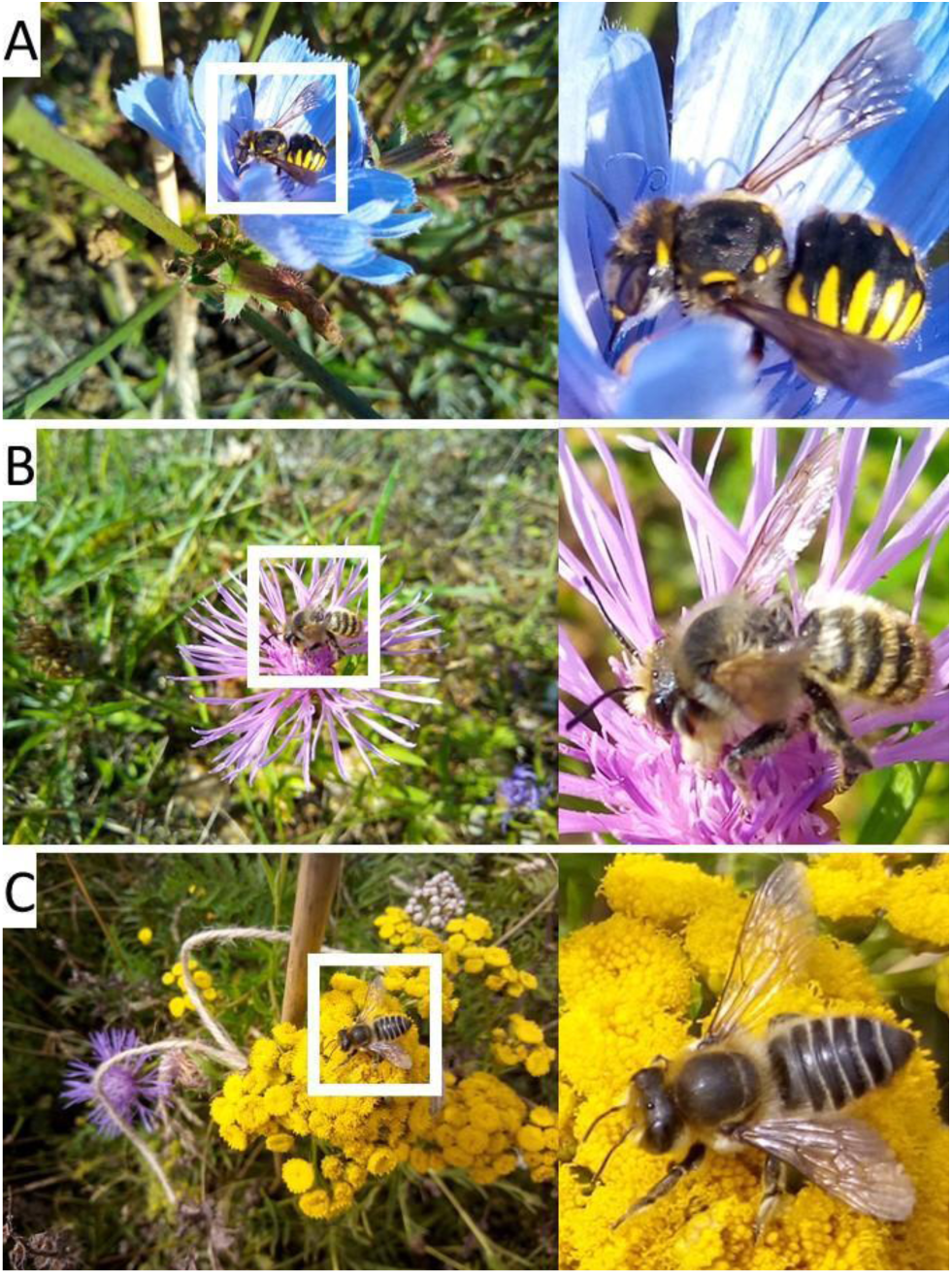
Examples of individuals from the Megachilidae family: (A) *Anthidium manicatum*, (B) showing abdominal position of *Osmia* genus, (C) showing abdominal position of *Megachile* genus.

However, the position of the abdomen when visiting flowers is a reliable feature that is easily seen in the images. In *Osmia*, the abdomen is generally horizontal or even curled under, whereas in *Megachile*, the end of the abdomen is typically angled upwards (Fig. 8B,C). There were four bounding boxes which were identified as individuals within the genus *Osmia*, and 149 which were identified as individuals within the genus *Megachile*. The remaining 21 bounding boxes could not be identified to the genus level; 14 of these were outside the ROI and the remaining seven were too blurry.

There were 181 bounding boxes which were identified to the family Melittidae. There are no unique traits that unite the family Melittidae, and thus all bounding boxes were assigned to one of two genera. The genus *Dasypoda*, to which three bounding boxes were assigned, are easily recognized by their conspicuously long hairs on the hind legs (Fig. 9A). Similarly, the genus *Macropis*, to which 178 bounding boxes were assigned, are recognized by a dense, light-coloured patch of long hairs on the hind tibiae (Fig. 9B). The identification of these occurrences as the genus *Macropis* was done cautiously, as image quality often does not provide a clear view of the scopa or wing venation. Identification to the species level in both genera requires close examination of the hairs on the legs and abdomen, which is not possible from our images.

**Figure 9.**
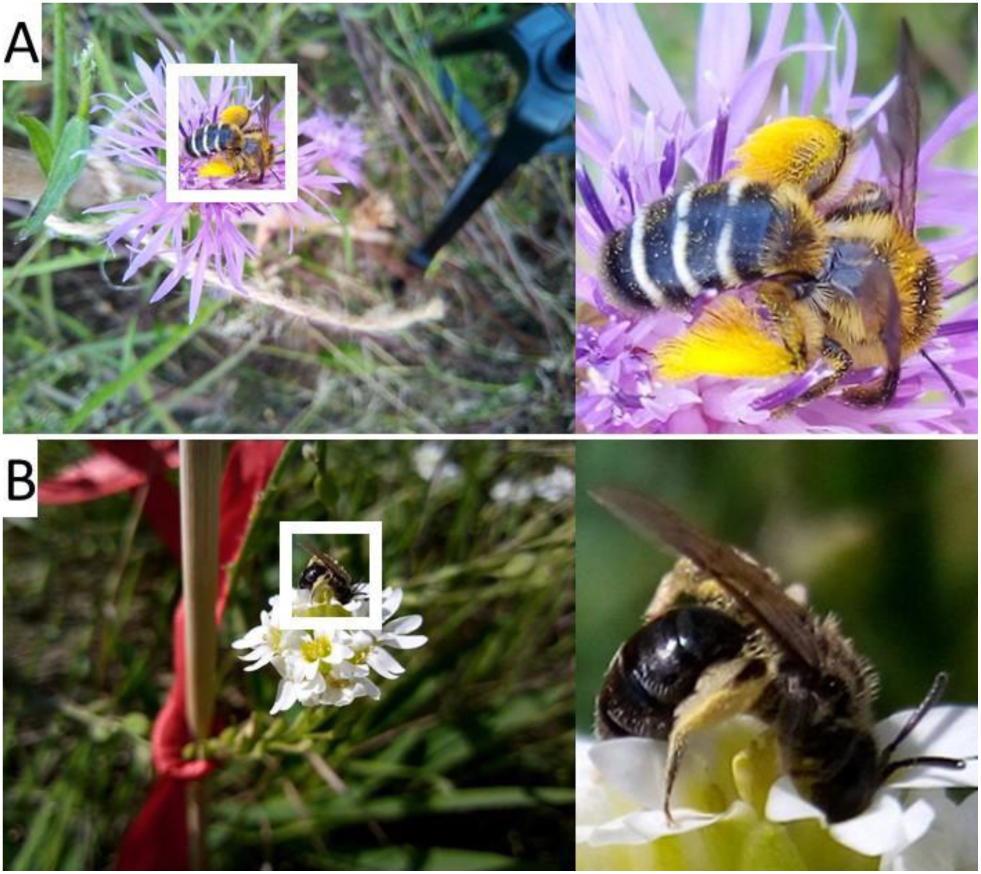
Examples of individuals from the Melittidae family: (A) of *Dasypoda* genus and its characteristically long leg hairs, (B) of *Macropis* genus and its characteristic tibial hairs.

Two larger wasps were also observed. One individual (five bounding boxes) belonged to the family Pompilidae and was a member of the genus *Episyron* due to the spines and reddish coloration of its legs. The other individual (two bounding boxes) belonged to the family Vespidae due to the position in which its wings were held at rest and its characteristic black and yellow coloration. We did not attempt to further identify this individual.

For the purposes of pollination ecology, it is desirable to identify insects to at least genus level, as this provides structural information on the plant-pollinator network that is similar to the species level (Rodrigues & Boscolo 2020). The ability of sampling with smartphones to capture images that enabled this level of identification was high. Of the 18,844 images identified to families within the order Hymenoptera, we did not attempt to identify 2,818 (15%) further because they are unlikely to be pollinators (families Cynipidae, Formicidae, Vespidae). Of those in which further identification was attempted, we identified 13,540 (71.9%) to genus. Of those that could not be identified to genus, 693 (3.7%) were outside the ROI, and thus would not have been the target for observations in pollination ecology (i.e., they are not visiting the focal flower we are observing). 171 (0.9%) were obscured, and thus even more expensive camera systems would not have enabled a closer identification. Finally, 1,612 (9%) were blurry. The blurry cases reflect the possible limitations of sampling with affordable smartphones, as more expensive camera systems might have better been able to capture sharper photos of small or fast-moving insects visiting the ROI.

### Identifying Diptera to lower taxonomic levels

Distinguishing between Diptera families requires clear images of characters of the wing veins in combination with other distinctive features such as body shape, patterning and/or colour. However, wing venation was not typically visible in our images. Therefore, identification to the family level posed significant challenges, and identification to the genus level proved even more demanding.

5,963 bounding boxes contained an individual in the order Diptera (Appendix Tab. V & Fig. 10). Of these, 3,066 (51.4%) could be identified to one of the six families/family-clusters: Anthomyiidae, Calliphoridae/Muscidae, Chyromyidae, Sarcophagidae/Tachinidae, Syrphidae and Tachinidae.

**Figure 10.**
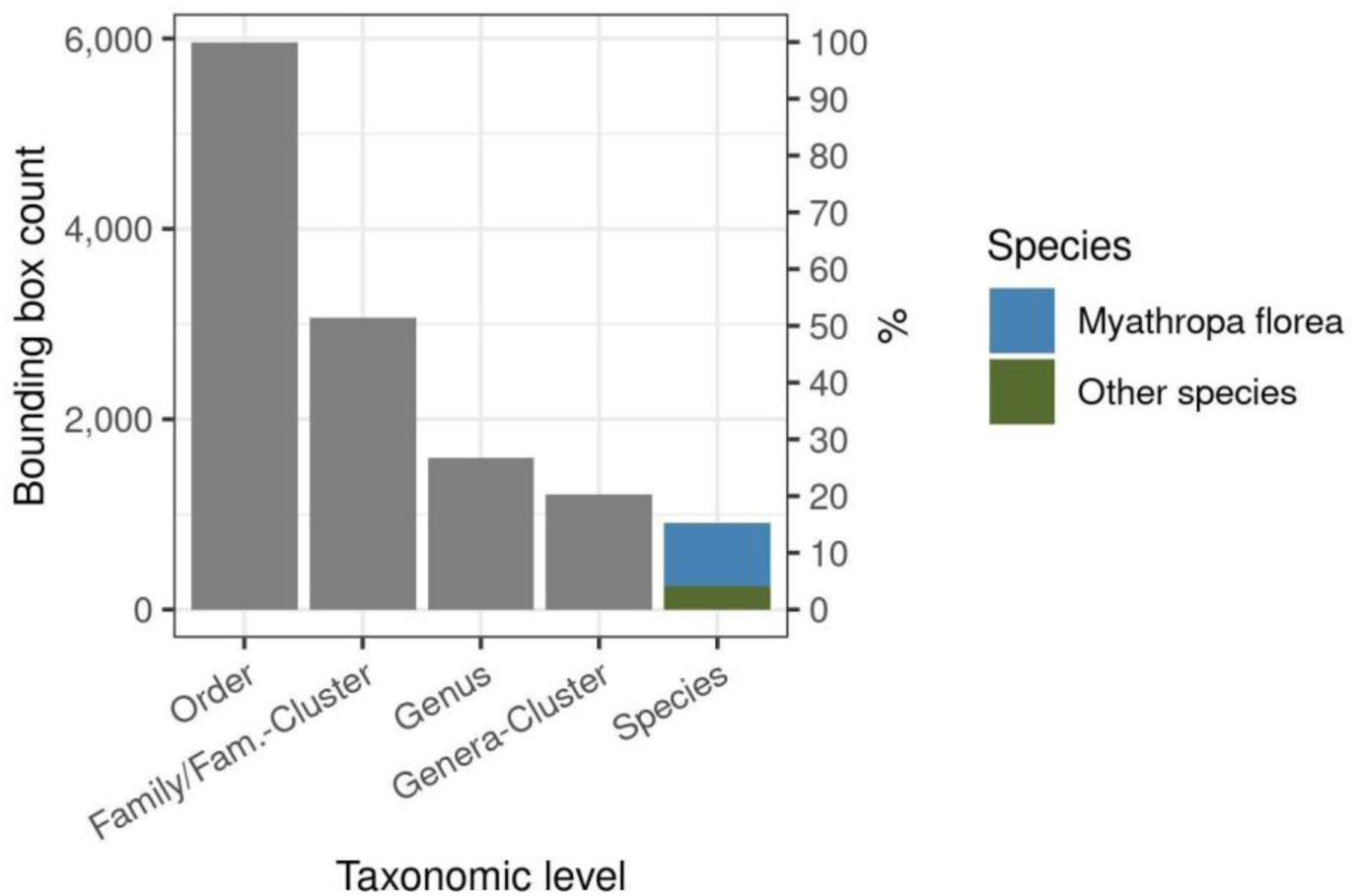
Bar plot illustrating the number of bounding boxes at different taxonomic levels for the flower visitors in the order Diptera. The primary y-axis (left) shows the count of boxes, while the secondary y-axis (right) displays the percentage of each taxonomic level relative to the total number of entries in the ‘Order’ level. This visualisation highlights the distribution and completeness of taxonomic information within the Diptera dataset. Note that *Myathropa florea* comprises 11.2% of the total bounding boxes labelled as Diptera (see also Appendix Tab. V)

The remaining 2,897 (48.6%) bounding boxes could not be identified to the family/family-cluster level (Fig. 11). Of these, in most cases the insect was too small and zooming in to see identifying features resulted in blurry, pixelated images. This occurred when we attempted to capture an entire inflorescence as a ROI, which resulted in blurry images of the small flies that visited flowers within that inflorescence (Fig. 11A). Similarly, if the phone was positioned too distant from the target flower, the images of small flies were blurry (Fig. 11B). Moreover, some of these flies were so tiny that identifying features could not be seen even when the camera was fairly close to the focal flower (e.g., Fig. 11C). We note that 697 of these cases were from one time-lapse photoset of one very tiny fly individual (Fig. 11C). Finally, 182 insects (6.3%) were outside the region of interest (ROI), so were not in the focus of the camera (Fig. 11D).

**Figure 11.**
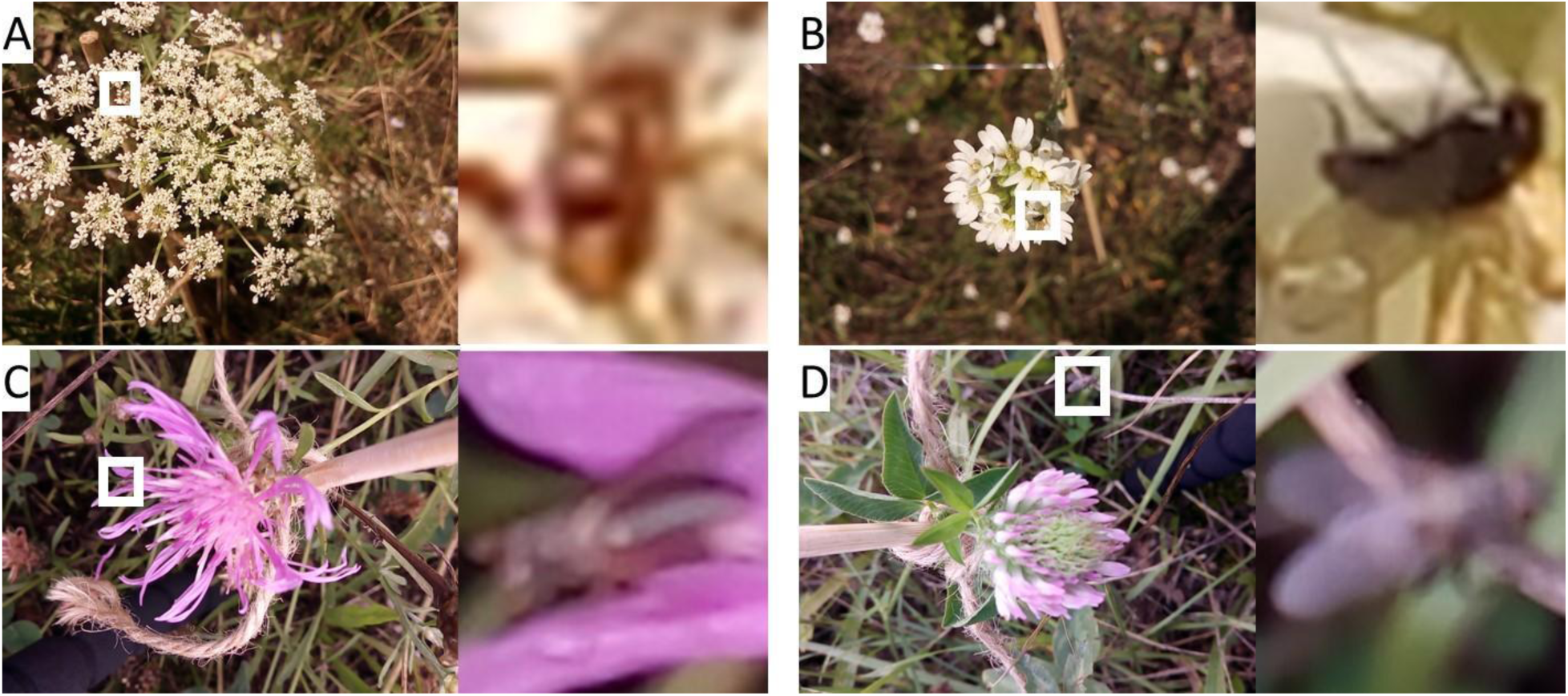
Examples of Diptera insects that could not be identified to the family or family-cluster taxonomic level: (A) the insect appears too small when attempting to capture the entire large inflorescence, (B) if the phone was too far from the flower, the insect appeared small and the identifying features are not visible, (C) Even if the phone is close to the flower and the insect is within the ROI, its features may not be visible due to its tiny size, (D) The insect is out of focus and outside the ROI.

Using a camera lens with more powerful optical zoom is a possible solution to these challenges, although it is important to note that there was a depth of field issue for such small flies. Increasing the optical zoom can further decrease the depth of field, making it even more challenging to maintain focus on small, moving insects. Although some identifying features may have been visible if the fly was at the correct focal depth, achieving this focus is difficult given the size of the flies.

We successfully identified all 17 bounding boxes in the family Anthomyiidae to genus *Anthomyia* due to their distinct characteristics in terms of size, shape, and markings. Further species-level identification is particularly challenging, even with the use of a microscope. Achieving species-level identification would necessitate clear views of the genitals to determine sex, as further identifying features are sex-dependant.

Despite originating from different superfamilies, we grouped the Calliphoridae and Muscidae families together due to the insects in our images exhibiting very similar characteristics, specifically silvery pilosity and metallic green-blue bodies. These traits, being the only ones visible in our images, allowed us to categorize 184 bounding boxes as belonging to the Calliphoridae-Muscidae group. However, this made reliable identification to one specific family challenging. Distinguishing between these two families would require high resolution photos of the face, thorax bristles and the wing veins. Similarly, identification to lower taxonomic levels within these families was not possible because the required identifying features (including body and facial bristles and hairs, wing veins, and antennae hairs) cannot be seen in the images.

A total of 57 bounding boxes were identified to the family Chyromyidae because of their small size, pale yellow body colour, clear wings and distinctive body shape. However, this family remains poorly understood, is underrepresented in collections, and species descriptions are only available for a subset of genera. Species identification within this family relies on features not seen in our images, such as thorax bristles, leg colour, body colour variations, leg bristles, and bristle colour.

Because of their similar appearances, it is challenging to differentiate between genera from the families Sarcophagidae and Tachinidae with similar wing veins, thick bristles and a black/silver checkerboard pattern on the abdomen. To address this issue, we created a separate family-cluster called “Sarcophagidae/Tachinidae”, to which 47 bounding boxes were assigned. The clear view of wing veins allowed us to further identify 32 of these bounding boxes (of a single insect) as either genus *Eliozeta* or *Clytiomya* within the family Tachinidae due to its distinct yellow markings. Distinguishing between *Eliozeta* and *Clytiomya* would require a microscope to see features including genitalia, and details in abdominal and thorax bristle arrangement.

There were 2,761 (46.3%) bounding boxes identified as Syrphidae family. Syrphidae can be more easily distinguished due to their diverse and unique shapes, patterns, and colours. For the family Syrphidae, 914 boxes could be identified to species: 224 *Episyrphus balteatus* (due to distinctive black and yellow “moustache” shape patterning on abdomen), 11 *Mesembrius pereginus* (due to distinctive thorax/abdomen patterning and the fact that there is a single species in Europe), 671 *Myathropa florea* (due to distinctive bright yellow colour and black patterning on abdomen, grey pattern on thorax, and wing veins; there is only one mainland European species), four *Scaeva pyrasti* (due to larger size, shining thorax, whiter downward-angled lunules/abdominal markings) and four *Syritta pipiens* (due to swollen hind femurs, colour pattern and general shape) (Fig. 12). It is worth noting that *Mesembrius pereginus* is a rare species, classified as threatened with extinction in Germany (Ssymank et al. 2011).

**Figure 12.**
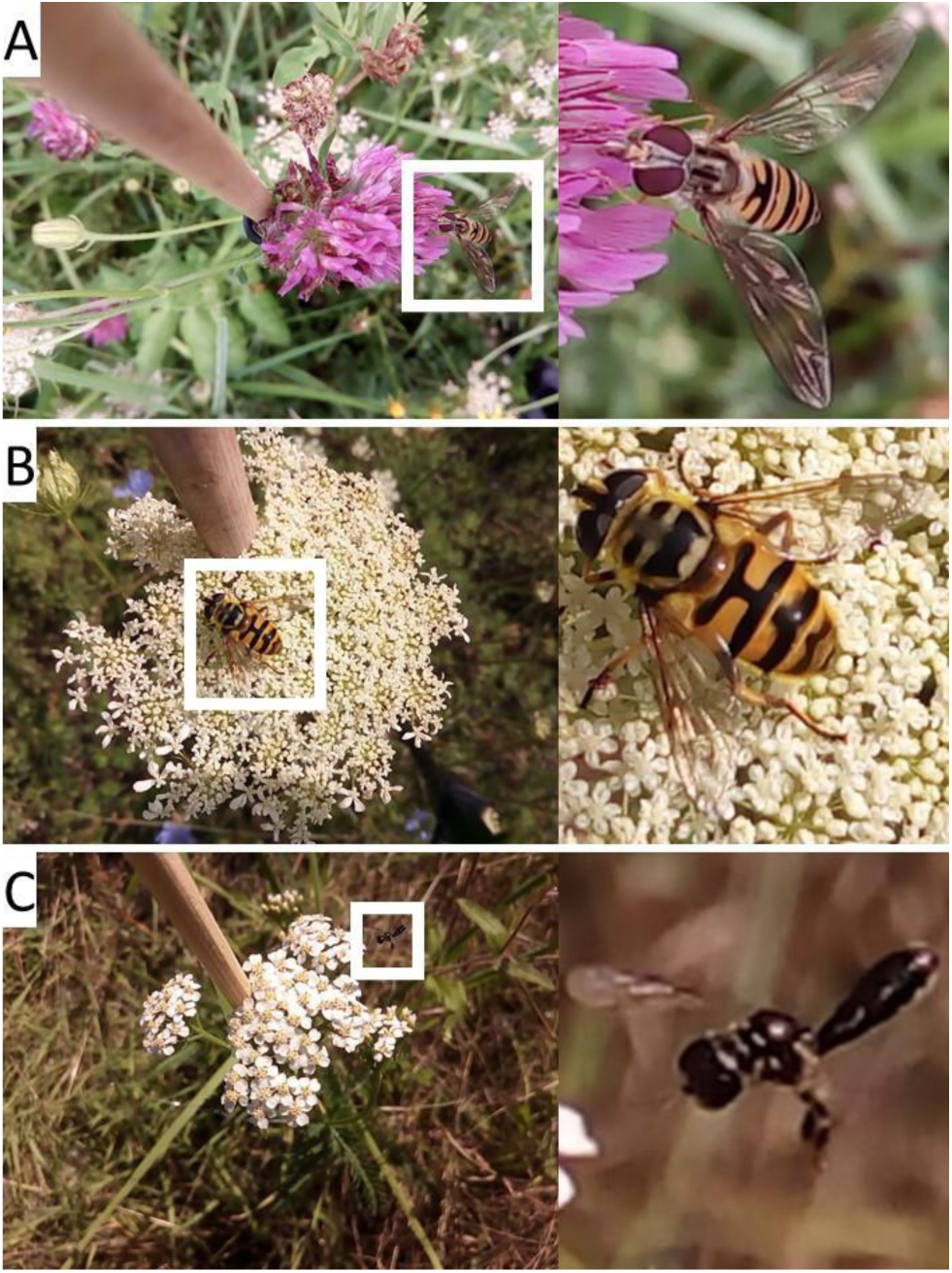
Examples of Syrphidae identified to species level: (A) *Episyrphus balteatus*, (B) *Myathropa florea*, (C) *Syritta pipiens*.

1,206 bounding boxes were identified to Syrphid genera groups. There are 467 species of Syrphidae that occur in the country of Germany (Ssymank et al. 2011). We created genera groups for these 467 Syrphid species (see details in Appendix Tab. VI), as there are many instances of genera bearing superficial resemblance to each other, requiring views of specific distinguishing features that would most likely not be visible in photos. The proposed groups vary in size, with the “wasp_mimic_banded” group having four species across two genera and the “small_black” group having 143 species across 15 genera.

There were only seven bounding boxes marked as Syrphidae that couldn’t be identified to the genera group level - two were outside the ROI, three displayed incomplete views of the insect (obscured), and two featured insects in flight (too blurry).

There were 341 bounding boxes identified as the genus *Eristalis*. This was possible due to their stout shape, large eyes, well-defined loop in the R4/5 wing vein, the forward and downward-projecting face and particular yellow abdomen markings. However, further identification to the species level is not possible due to the natural and seasonal variation in coloration and markings between sexes within a species, as well as morphological overlap between different sexes across species. A microscope would be required to distinguish between *Eristalis* species, to show details such as hairs on wing calypter, body and face hair colour, leg colour, facial stripe view, antennal hairs, wing veins, and male genitalia.

Four bounding boxes were identified as the genus *Pipiza* due to the shape, body hair colour, leg colour and darkened wing edge. These small black flies can only be classified to the genus level with our current photos. Distinguishing between species requires details such as antennal placement, eye angle, facial features, leg hairs and various other characteristics related to size, colour, and patterning, which are not discernible in our images.

There were 289 bounding boxes identified as *Sphaerophoria*, a genus which is also part of the genera cluster “black yellow elongated”. The *Sphaerophoria* genus can be more easily identified from other genera in the cluster if the yellow stripe on thorax sides is visible. Males of *S. scripta* might be identifiable to species level if the abdomen is shown to be longer than the wings, but the photo angle plays a crucial role.

## Lessons learned and ideas for future development

Batteries: We did not experience any battery issues in the field. We used the phones for several hours each day, and we ensured that they were recharged at the end of each day.

Heat and humidity: Notably, some of our smartphones shut down during extreme heat. Similarly, it has been reported that there is also a risk that they might not charge under high humidity conditions (pers. comm. Robert Tropek, Charles University, Prague). To mitigate the heat issue, we used white casings that reflect light (a simple solution we employed involved using a piece of paper). This issue may also arise in microcomputers, where heatsinks and coolers are essential to maintain the device at operational temperatures (e.g., Sittinger et al. 2023). Nevertheless, surveillance cameras designed for outdoor use appear to be more resilient to such conditions, though they are typically more expensive than smartphones (pers. comm. Robert Tropek, Charles University, Prague).

Image resolution: While it is tempting to increase the image resolution to the maximum offered by the smartphone model, it is not advisable because storing a larger amount of data can become impractical for data transfer and logistics without providing additional taxonomic details, particularly if the focus is not sharp enough. Most images in our dataset were taken at a custom resolution of 1600 x 1200 pixels. Generally, a 1200 x 1200 resolution is sufficient for capturing essential insect traits, provided the insect is in focus.

Open-source time-lapse apps: The Open Camera app provides various custom features essential for our fieldwork, such as custom resolution, manual focus, custom time steps, automatic exposure, file format, image compression, and custom image folder naming. However, despite its versatility, Open Camera lacks the ability to define a rectangular region of interest (ROI). We were unable to find any open-source app for Android OS that combines this feature with the other requirements.

Predefined ROI: There is the necessity for an app feature that can capture images based on a predefined rectangular region of interest (ROI) that can be set at the beginning of the time-lapse session. This ROI should be adjusted to frame only the target flower or inflorescence, possibly including a buffer to account for wind movements, and minimise background noise. Establishing such a ROI ensures that the camera focuses on visiting insects, while also maximising their occupancy in the image, thereby facilitating identification.

Dual lenses to capture insects in the ROI: Employing two distinct lenses could be advantageous: one for capturing close-up images, providing greater detail for smaller insects (e.g., smaller than 2 cm); and another with a field of view or region of interest (ROI) configured to accommodate larger insects (larger than 2 cm, such as sizable Lepidoptera). This approach can help ensure comprehensive image collection across a range of insect sizes.

Furthermore, when selecting a smartphone for pollinator monitoring, it is recommended to prioritise larger optical magnification (i.e., longer focal lengths) over high resolution sensors (e.g., a usual 12 MP camera sensor is more than enough). The use of recent smartphone models, which are often equipped with 2X optical zoom lenses, is suggested. Digital zoom is not advisable due to the resulting information loss. An optical zoom allows for a reduction in background noise in the frame without the need to get too close to the flower, which may disturb the visitors.

Lack of butterfly observations: Few butterflies were observed in our study, potentially due to factors such as time of day, or equipment disturbance.

Data transfer: Data transfer directly from phones via USB cables was slow, prone to interruptions, and carries a risk of data loss. Such loss can occur when users mistakenly believe the data has already been downloaded and proceed to delete it from the phone to prepare for the next day. However, this issue is not unique to smartphones and can affect any camera system, unless live streaming to a server is utilised where internet coverage is available in the field. To expedite data transfer, an alternative approach involved removing the micro SD card from the smartphone and uploading the data directly using a computer’s microSD card reader. However, this method necessitated frequent opening and closing of the phone, increasing the risk of breaking the protective casing or causing other damage due to the repeated exposure of fragile components to sharp tools. More recent phone models might allow easier access to the SD cards and/or faster download speeds.

Image capture speed: Affordable smartphones are generally slower at capturing images compared to devices like Raspberry Pi units. Although we configured the Open Camera app to take an image every second, in practice, due to overheating or processing power limitations, smartphones struggle to achieve this rate, typically capturing one image every 1.5 to 2 seconds. In contrast, microcomputers can rapidly capture dozens of images per second (e.g., Droissart et al. 2021; Sittinger et al. 2023) without requiring significant processing power. Nonetheless, even at reduced frame rates of 1.5 or 2 seconds, we were typically able to capture multiple images of each visiting individual, enabling taxonomists to identify insects by observing them from various angles. Additionally, it is worth noting that reducing the time step to capture several frames per second would quickly exhaust the storage capacity of the micro SD cards on the smartphones. For example, with an image resolution of 1200 x 1200 pixels and assuming a JPG file format that allows 1 MB per frame, capturing at 2 frames per second could easily generate between 7 and 8 GB in just one hour.

Many images without insects in them: Due to the time-lapse approach employed in capturing images, 93% of the images in our dataset did not contain insects. It is customary to observe such results with time-lapse camera systems; for instance, Ruczyński et al. (2020), noted that over 90% of their captured photographs were devoid of insects.

Ideally, a custom trigger for pollinators would help reduce the volume of stored data. However, several challenges exist, such as insects not emitting heat, often being too small to trigger motion sensors, or sensors being activated by wind movement. There are ongoing efforts to develop AI triggers for pollinators, which involve real-time AI pipelines (edge AI) powered by nano GPUs in the field (e.g., Bjerge et al. 2022; Sittinger et al. 2023).

Unfortunately, smartphones do not come yet with GPUs like those found in Jetson Nano from NVIDIA (2019) or Coral microcomputers from Google (Coral 2020), which allow for edge AI (rapid insect detection on the device). However, fast on-device detection carries significant risks, as it may disadvantage small species or any taxa that the AI was not trained on. For example, Van Horn et al. (2018) found that in their iNaturalist dataset, small organisms posed a challenge for classification models, even when they were accurately localised by the AI architecture. van Klink et al. (2022) underlined that AI struggles with rare species due to limited data and tends to overpredict common species with abundant data. Consequently, edge AI, despite its potential advantages, could pose a significant bottleneck and exhibit bias towards larger and more well-represented species. Another solution might be to process images either on the smartphone, running an AI model in the background and not in real-time (though this is energy-intensive and may not be feasible) or upload images in real time to a GPU server, which offers superior detection and classification capabilities (though this requires good internet coverage in the field).

Despite the high percentage of images without insects, it’s important to recognize the value of this seemingly ‘empty’ data. Particularly, an edge AI device that records only detected insects may retain biases, as a human annotator will not see the ‘empty’ data to identify potential misses, which could be used for retraining and updating the AI model.

Our findings demonstrate that, despite certain limitations, smartphones can be an effective tool for pollinator monitoring when researchers take appropriate measures to optimise their usage. Nearly all insects can be identified at the order level. Hymenopterans that are known to pollinate can often be identified to the genus level. Dipteran identification works best when there are sharp photos of the wings, which was often not achieved with the smartphones, and thus Dipterans could not be identified as well from images taken with smartphones as Hymenopterans. As technology continues to evolve, smartphones are becoming increasingly accessible, affordable and user-friendly, making them an attractive option for researchers aiming to monitor pollinators.

## APPENDICES

Additional supporting information may be found in the online version of this article:

Appendix Table I. Gear details and costs.

Appendix Table II. Sampled plant species and number of annotated images.

Appendix Table III. Hymenoptera - counts of bounding boxes.

Appendix Table IV. Bombus species grouped in morpho-species.

Appendix Table V. Diptera - counts of bounding boxes.

Appendix Table VI. Syrphid species grouped in clusters.

Appendix Table VII. Fieldwork protocol.

Appendix Table VII. List of sites

Appendix Figure I. Frequency of images by hour of the day.

## ACKNOWLEDGEMENTS

The author thanks Bilyana Stoykova, Ricardo Urrego Alvarez, Emil Cyranka, Anna Scheiper for helping with field work and manually placing the bounding boxes and Anne-Kathrin Thomas for logistical support. We are grateful to Amibeth Thompson for providing metadata for the sites. This research was funded by the Helmholtz AI initiative (Information & Data Science) Pollination Artificial Intelligence (ZT-I-PF-5-115), led by Prof. Tiffany M. Knight and Prof. Hannes Taubenboeck, the Helmholtz Recruitment Initiative of the Helmholtz Association to Tiffany M. Knight, and iDiv (German Research Foundation FZT 118).

## DATA AVAILABILITY STATEMENT

The image dataset associated with this research is available from the corresponding author upon reasonable request, owing to its substantial storage size. The remaining data for this study are published as supplementary material alongside the online version of this article. The image annotation data and the R code used for creating some of the figures and tables in this manuscript and appendices are available in our GitHub repository at: https://github.com/valentinitnelav/pollinator-image-annotation

