## Appendices for "Utilising affordable smartphones and open-source time-lapse photography for monitoring pollinators"

**Appendix Figure I. Frequency of images by hour of the day**

Distribution of the number of images by hour of the day for the 213 annotated "plant folders" (each representing approximately 1 hour of time-lapse images). The red dashed lines represent 95% quantile-based confidence intervals, indicating the range within which 95% of the images fall in the distribution.

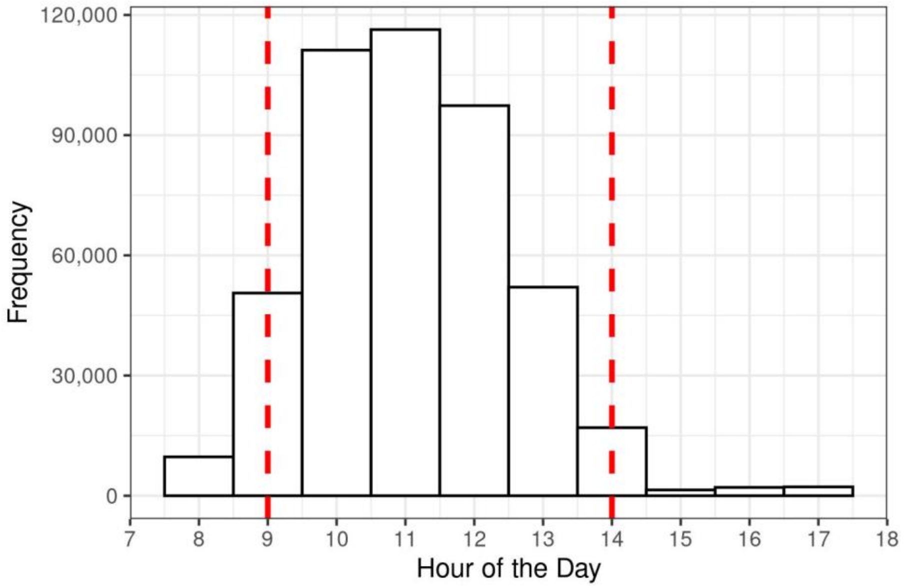

**Appendix Table I. Gear details and costs**

| ID | Product | Nr. units | URL (last checked 2023-11-30) | Unit price (€, 2021) |
|---|---|---|---|---|
| 1 | Blackview A60 smartphone | 6 | https://www.blackview.hk/products/item/a60 | 85 |
| 2 | Oukitel WP17 outdoor smartphone | 2 | https://oukitel.com/products/oukitel-wp17-rugged-phone | 130 |
| 3 | Djroll 36000 mAh Qi wireless solar power bank | 2 | https://www.amazon.de/dp/B08P428WK5 | 40 |
| 4 | Sweye solar power bank 26800 mAh | 3 | https://www.amazon.de/dp/B08CV3JV7C | 22 |
| 5 | Hermitshell Poweradd EnergyCell 10000 mAh | 3 | https://www.amazon.de/dp/B07T8N2B29 | 11 |
| 6 | Everesta aluminium mobile phone tripod | 8 | https://www.amazon.de/dp/B0725GDDQX | 23 |
| 7 | SanDisk micro SD card, 32 Gb | 8 | https://www.amazon.de/dp/B06XWMQ81P | 9 |

Example of gear combination (total cost per unit):
Product id 1+5+6+7 results in 85+11+23+9 = 128 EUR (low cost option; prices in 2021)
Product id 2+3+6+7 results in 130+40+23+9 = 202 EUR (most expensive option; prices in 2021)

**Appendix Table II. List of sampled plant species and number of annotated images.**

"Nr. folders": Number of one-hour folders containing time-lapse images (proxy for the number of hours spent observing each plant species);
"Nr. img.": Total number of images visually inspected for the presence of insects;
"Nr. img. w. insect": Number of images containing a insect;
"% img w. insect": Percentage of images containing an insect, relative to the total number of images.

| Id. | Plant | Nr. folders | Nr. img. | Nr. img. w. insect | % img w. insect |
|---|---|---|---|---|---|
| 1 | *Achillea millefolium* | 4 | 9,154 | 1,204 | 13.15 |
| 2 | *Asteraceae* "white" | 1 | 1,832 | 325 | 17.74 |
| 3 | *Berteroa incana* | 5 | 13,100 | 335 | 2.56 |
| 4 | *Bunias orientalis* | 3 | 8,328 | 376 | 4.51 |
| 5 | *Carduus acanthoides* | 10 | 22,040 | 1,776 | 8.06 |
| 6 | *Centaurea jacea* | 54 | 118,476 | 11,413 | 9.63 |
| 7 | *Centaurea scabiosa* | 3 | 3,814 | 681 | 17.86 |
| 8 | *Centaurea stoebe* | 3 | 8,308 | 233 | 2.8 |
| 9 | *Cichorium intybus* | 28 | 61,348 | 1,879 | 3.06 |
| 10 | *Cirsium vulgare* | 2 | 3,748 | 514 | 13.71 |
| 11 | *Clematis vitalba* | 7 | 13,798 | 2,072 | 15.02 |
| 12 | *Crepis biennis* | 11 | 21,224 | 832 | 3.92 |
| 13 | *Daucus carota* | 15 | 31,054 | 4,713 | 15.18 |
| 14 | *Echium vulgare* | 3 | 6,078 | 53 | 0.87 |
| 15 | *Erigeron annuus* | 2 | 3,243 | 191 | 5.89 |
| 16 | *Hypericum perforatum* | 6 | 10,049 | 255 | 2.54 |
| 17 | *Hypochaeris radicata* | 2 | 3,986 | 91 | 2.28 |
| 18 | *Lamium purpureum* | 1 | 2,505 | 13 | 0.52 |
| 19 | *Leucanthemum vulgare* | 1 | 2,577 | 838 | 32.52 |
| 20 | *Lotus corniculatus* | 1 | 1,988 | 57 | 2.87 |
| 21 | *Melilotus albus* | 2 | 4,312 | 86 | 1.99 |

| | | | | | |
|---|---|---|---|---|---|
| 22 | *Origanum vulgare* | 3 | 7,099 | 437 | 6.16 |
| 23 | *Picris hieracioides* | 17 | 36,037 | 1,172 | 3.25 |
| 24 | *Scorzoneroides autumnalis* | 1 | 3,802 | 69 | 1.81 |
| 25 | *Sedum sp* | 1 | 1,828 | 14 | 0.77 |
| 26 | *Senecio inaequidens* | 2 | 4,394 | 65 | 1.48 |
| 27 | *Silene latifolia* | 1 | 3,037 | 5 | 0.16 |
| 28 | *Tanacetum vulgare* | 6 | 12,729 | 2,054 | 16.14 |
| 29 | *Trifolium medium* | 1 | 3,903 | 23 | 0.59 |
| 30 | *Trifolium pratense* | 12 | 26,083 | 1,416 | 5.43 |
| 31 | *Trifolium repens* | 1 | 1,907 | 6 | 0.31 |
| 32 | *Verbascum densiflorum* | 1 | 2,116 | 144 | 6.81 |
| 33 | *Vicia villosa* | 3 | 6,159 | 153 | 2.48 |
| | **TOTAL** | **213** | **460,056** | **33,495** | **7.28** |

**Appendix Table III. Counts of bounding boxes containing a flower visitor in the order Hymenoptera**

| Id. | Family | Genus | Morphospecies | Species | Nr. boxes | % boxes |
|---|---|---|---|---|---|---|
| 1 | NA | | | | 2,138 | 10.187 |
| 2 | Andrenidae | | | | 355 | 1.692 |
| 3 | Andrenidae | *Andrena* | | | 253 | 1.206 |
| 4 | Apidae | | | | 7 | 0.033 |
| 5 | Apidae | *Anthophora* | | | 1 | 0.005 |
| 6 | Apidae | *Apis* | | *mellifera* | 3,307 | 15.757 |
| 7 | Apidae | *Bombus* | | | 115 | 0.548 |
| 8 | Apidae | *Bombus* | black | | 7 | 0.033 |
| 9 | Apidae | *Bombus* | red_tailed | | 3,669 | 17.482 |
| 10 | Apidae | *Bombus* | red_tailed | *lapidarius* | 7 | 0.033 |
| 11 | Apidae | *Bombus* | red_yellow | | 698 | 3.326 |
| 12 | Apidae | *Bombus* | striped | | 10 | 0.048 |
| 13 | Apidae | *Bombus* | white_tailed | | 672 | 3.202 |
| 14 | Colletidae | *Colletes* | | | 1 | 0.005 |
| 15 | Colletidae | *Colletes* | | *cunicularius* | 1 | 0.005 |
| 16 | Colletidae | *Hylaeus* | | | 402 | 1.915 |
| 17 | Cynipidae | | | | 390 | 1.858 |
| 18 | Formicidae | | | | 2,426 | 11.560 |
| 19 | Halictidae | | | | 2,092 | 9.968 |
| 20 | Halictidae | *Halictus* | | | 2,250 | 10.721 |
| 21 | Halictidae | *Halictus* | | *scabiosae* | 287 | 1.368 |
| 22 | Halictidae | *Halictus* | | *subauratus* | 422 | 2.011 |
| 23 | Halictidae | *Lasioglossum* | | | 941 | 4.484 |
| 24 | Halictidae | *Lasioglossum* | | *calceatum* | 28 | 0.133 |
| 25 | Halictidae | *Sphecodes* | | | 133 | 0.634 |
| 26 | Megachilidae | | | | 21 | 0.100 |
| 27 | Megachilidae | *Anthidium* | | *manicatum* | 13 | 0.062 |
| 28 | Megachilidae | *Megachile* | | | 149 | 0.710 |
| 29 | Megachilidae | *Osmia* | | | 4 | 0.019 |
| 30 | Melittidae | *Dasypoda* | | | 3 | 0.014 |
| 31 | Melittidae | *Macropis* | | | 178 | 0.848 |

| | | | | | | |
|---|---|---|---|---|---|---|
| 32 | Pompilidae | *Episyron* | | | 5 | 0.024 |
| 33 | Vespidae | | | | 2 | 0.010 |
| | **TOTAL** | | | | **20,987** | **100** |

**Appendix Table IV. List of Hymenoptera species in the *Bombus* genera grouped in morpho-species based on observable traits in smartphone images.**

These species can occur in Germany and bordering countries. The meanings of the abbreviations, columns and additional helpful notes associated with this table are listed at its end.

| Species | Sex | Morpho-species | Anterior thoracic colour | Central thoracic colour | Posterior thoracic colour | T1 colour | T2 colour | T3 colour | T4 colour | T5+ colour |
|---|---|---|---|---|---|---|---|---|---|---|
| *argillaceus* | M | white_tailed | y | b | y | y | b | b | w | w |
| *argillaceus* | F | white_tailed | y | b | y | y | b | b | w | w |
| *argillaceus* | Q | black_tailed | y | b | y | b | b | b | b | b |
| *armeniacus* | M | striped | w | b | w | y | y | y | y | y |
| *armeniacus* | F | striped | y | b | y | y | m | y | y | y |
| *armeniacus* | Q | striped | y | b | y | y | m | y | y | y |
| *balteatus* | M | striped | y | b | y | y | y | b | m | m |
| *balteatus* | F | striped | y | b | y | y | y | b | m | m |
| *balteatus* | Q | striped | y | b | y | y | y | b | m | m |
| *barbutellus* | M | white_tailed | y | b | y | m | b | m | w | w |
| *barbutellus* | Q | white_tailed | y | b | y | m | b | m | w | w |
| *bohemicus* | M | white_tailed | y | b | m | m | b | m | w | m |
| *bohemicus* | Q | white_tailed | y | b | b | b | b | m | w | m |
| *campestris* | M | striped | y | b | y | y | b | m | m | m |
| *campestris* | Q | striped | y | b | y | b | b | m | m | m |
| *cingulatus* | M | white_tailed | r | b | r | b | b | b | b | w |
| *cingulatus* | F | white_tailed | r | b | r | b | b | b | b | w |
| *cingulatus* | Q | white_tailed | r | b | r | b | b | b | b | w |
| *confusus* | M | red_tailed | b | b | b | b | b | b | r | r |
| *confusus* | F | red_tailed | b | b | b | b | b | b | r | r |
| *confusus* | Q | red_tailed | b | b | b | b | b | b | r | r |
| *confusus* | M | white_tailed | y | b | m | y | b | b | w | w |
| *confusus* | F | white_tailed | y | b | m | y | b | b | w | w |
| *confusus* | Q | white_tailed | y | b | m | y | b | b | w | w |
| *consobrinus* | M | red_yellow | r | r | r | r | r | m | w | w |
| *consobrinus* | F | red_yellow | r | r | r | r | r | m | w | w |
| *consobrinus* | Q | red_yellow | r | r | r | r | r | m | w | w |
| *cryptarum* | M | white_tailed | y | b | b | b | y | b | w | w |
| *cryptarum* | F | white_tailed | y | b | b | b | y | b | w | w |
| *cryptarum* | Q | white_tailed | y | b | b | b | y | b | w | w |
| *cullumanus* | M | red_tailed | y | b | y | y | y | b | r | r |
| *cullumanus* | F | red_tailed | b | b | b | b | b | b | r | r |
| *cullumanus* | Q | red_tailed | b | b | b | b | b | b | r | r |
| *cullumanus* | M | red_tailed | y | b | y | y | y | b | r | r |
| *cullumanus* | F | red_tailed | y | b | y | y | y | b | r | r |
| *cullumanus* | Q | red_tailed | y | b | y | y | y | b | r | r |
| *distinguendus* | M | striped | y | b | y | y | y | y | y | y |
| *distinguendus* | F | striped | y | b | y | y | y | y | y | y |
| *distinguendus* | Q | striped | y | b | y | y | y | y | y | y |
| *flavidus* | M | striped | y | b | m | m | b | y | y | y |

| | | | | | | | | | | |
|---|---|---|---|---|---|---|---|---|---|---|
| *flavidus* | Q | striped | y | b | m | m | m | m | r | r |
| *fragrans* | M | striped | y | b | y | y | y | y | y | y |
| *fragrans* | F | striped | y | b | y | y | y | y | y | y |
| *fragrans* | Q | striped | y | b | y | y | y | y | y | y |
| *gerstaeckeri* | M | white_tailed | y | y | y | y | b | b | w | w |
| *gerstaeckeri* | F | white_tailed | y | y | y | y | b | b | w | w |
| *gerstaeckeri* | Q | white_tailed | y | y | y | y | b | b | w | w |
| *haematurus* | M | striped | y | b | m | m | y | y | b | b |
| *haematurus* | F | striped | y | b | b | b | y | y | b | b |
| *haematurus* | Q | striped | y | b | b | b | y | y | b | b |
| *hortorum* | M | white_tailed | y | b | y | y | b | b | w | w |
| *hortorum* | F | white_tailed | y | b | y | y | b | b | w | w |
| *hortorum* | Q | white_tailed | y | b | y | y | b | b | w | w |
| *hortorum* | M | black | b | b | b | b | b | b | b | b |
| *hortorum* | F | black | b | b | b | b | b | b | b | b |
| *hortorum* | Q | black | b | b | b | b | b | b | b | b |
| *humilis* | M | red_yellow | m | m | m | m | m | m | m | m |
| *humilis* | F | red_yellow | m | m | m | m | m | m | m | m |
| *humilis* | Q | red_yellow | m | m | m | m | m | m | m | m |
| *hyperboreus* | M | black_tailed | y | b | y | y | y | b | b | b |
| *hyperboreus* | F | black_tailed | y | b | y | y | y | b | b | b |
| *hyperboreus* | Q | black_tailed | y | b | y | y | y | b | b | b |
| *hypnorum* | M | red_thorax | r | r | r | r | b | b | m | w |
| *hypnorum* | F | red_thorax | r | r | r | b | b | b | m | w |
| *hypnorum* | Q | red_thorax | r | r | r | b | b | b | m | w |
| *jonellus* | M | white_tailed | y | b | y | y | m | b | b | w |
| *jonellus* | F | white_tailed | y | b | y | y | b | b | w | w |
| *jonellus* | Q | white_tailed | y | b | y | y | b | b | w | w |
| *laesus* | M | primarily yellow | y | r | y | y | y | y | y | y |
| *laesus* | F | primarily yellow | y | r | y | y | y | y | y | y |
| *laesus* | Q | primarily yellow | y | r | y | y | y | y | y | y |
| *lapidarius* | M | red_tailed | y | b | m | b | b | b | r | r |
| *lapidarius* | F | red_tailed | b | b | b | b | b | b | r | r |
| *lapidarius* | Q | red_tailed | b | b | b | b | b | b | r | r |
| *lapponicus* | M | red_tailed | y | b | m | y | r | r | y | y |
| *lapponicus* | F | red_tailed | m | b | m | m | r | r | r | r |
| *lapponicus* | Q | red_tailed | m | b | m | m | r | r | r | r |
| *lucorum* | M | white_tailed | y | b | y | b | y | b | w | w |
| *lucorum* | F | white_tailed | y | b | b | b | y | b | w | w |
| *lucorum* | Q | white_tailed | y | b | b | b | y | b | w | w |
| *magnus* | M | white_tailed | y | b | b | b | y | b | w | w |
| *magnus* | F | white_tailed | y | b | b | b | y | b | w | w |
| *magnus* | Q | white_tailed | y | b | b | b | y | b | w | w |
| *mesomelas* | M | striped | w | b | w | m | w | w | y | y |
| *mesomelas* | F | striped | y | b | y | y | r | y | y | y |
| *mesomelas* | Q | striped | y | b | y | y | r | y | y | y |
| *monticola* | M | red_tailed | y | b | m | m | r | r | r | r |
| *monticola* | F | red_tailed | y | b | m | b | m | r | r | r |
| *monticola* | Q | red_tailed | y | b | m | b | r | r | r | r |
| *mucidus* | M | striped | y | b | y | y | b | m | m | m |
| *mucidus* | F | striped | y | b | y | y | b | m | m | m |
| *mucidus* | Q | striped | y | b | y | y | b | m | m | m |
| *muscorum* | M | red_yellow | r | r | r | y | y | y | y | y |
| *muscorum* | F | red_yellow | r | r | r | y | y | y | y | y |
| *muscorum* | Q | red_yellow | r | r | r | y | y | y | y | y |

| | | | | | | | | | | |
|---|---|---|---|---|---|---|---|---|---|---|
| *norvegicus* | M | white_tailed | y | b | b | m | b | w | w | m |
| *norvegicus* | Q | white_tailed | y | b | b | b | b | w | w | w |
| *pascuorum* | M | red_yellow | r | r | r | y | m | m | r | r |
| *pascuorum* | F | red_yellow | r | r | r | y | m | m | r | r |
| *pascuorum* | Q | red_yellow | r | r | r | y | m | m | r | r |
| *polaris* | M | striped | y | m | y | y | y | m | m | y |
| *polaris* | F | black_tailed | y | b | y | y | y | y | b | b |
| *polaris* | Q | black_tailed | y | b | y | y | y | m | b | b |
| *pomorum* | M | red_tailed | b | b | b | w | w | r | r | r |
| *pomorum* | F | red_tailed | b | b | b | m | m | r | r | r |
| *pomorum* | Q | red_tailed | b | b | b | m | m | r | r | r |
| *pratorum* | M | red_tailed | y | b | m | y | y | b | r | r |
| *pratorum* | F | red_tailed | y | b | b | b | m | b | r | r |
| *pratorum* | Q | red_tailed | y | b | b | b | m | b | r | r |
| *pyrenaeus* | M | red_tailed | y | b | y | y | m | b | r | r |
| *pyrenaeus* | F | red_tailed | y | b | y | y | m | b | r | r |
| *pyrenaeus* | Q | red_tailed | y | b | y | y | m | b | r | r |
| *quadricolor* | M | white_tailed | y | b | b | y | b | b | w | w |
| *quadricolor* | Q | white_tailed | y | b | b | b | b | m | w | y |
| *ruderarius* | M | red_tailed | b | b | b | b | b | b | r | r |
| *ruderarius* | F | red_tailed | b | b | b | b | b | b | r | r |
| *ruderarius* | Q | red_tailed | b | b | b | b | b | b | r | r |
| *ruderatus* | M | white_tailed | y | b | y | y | b | m | w | w |
| *ruderatus* | F | white_tailed | y | b | y | y | b | m | w | w |
| *ruderatus* | Q | white_tailed | y | b | y | y | b | m | w | w |
| *ruderatus* | M | black | b | b | b | b | b | b | b | b |
| *ruderatus* | F | black | b | b | b | b | b | b | b | b |
| *ruderatus* | Q | black | b | b | b | b | b | b | b | b |
| *rupestris* | M | red_tailed | m | b | m | m | m | m | r | r |
| *rupestris* | Q | red_tailed | m | b | b | b | b | b | r | r |
| *schrencki* | M | red_yellow | r | r | r | r | r | y | y | y |
| *schrencki* | F | red_yellow | r | r | r | y | r | y | y | y |
| *schrencki* | Q | red_yellow | r | r | r | y | r | y | y | y |
| *semenoviellus* | M | white_tailed | y | b | y | m | m | b | w | w |
| *semenoviellus* | F | white_tailed | y | b | y | m | m | b | w | w |
| *semenoviellus* | Q | white_tailed | y | b | y | m | m | b | w | w |
| *sichelii* | M | striped | y | b | y | y | b | b | y | y |
| *sichelii* | F | striped | y | b | y | y | b | b | y | y |
| *sichelii* | Q | striped | y | b | y | y | b | b | y | y |
| *soroeensis* | M | white_tailed | y | b | b | y | y | b | y | w |
| *soroeensis* | F | white_tailed | y | b | b | b | y | b | y | w |
| *soroeensis* | Q | white_tailed | y | b | b | b | y | b | y | w |
| *soroeensis* | M | red_tailed | y | b | b | y | b | b | r | r |
| *soroeensis* | F | red_tailed | b | b | b | b | b | b | r | r |
| *soroeensis* | Q | red_tailed | b | b | b | b | b | b | r | r |
| *sporadicus* | M | white_tailed | y | b | y | y | y | b | w | w |
| *sporadicus* | F | white_tailed | y | b | y | y | y | b | w | w |
| *sporadicus* | Q | white_tailed | y | b | y | y | y | b | w | w |
| *subterraneus* | M | striped | y | b | y | y | b | m | m | m |
| *subterraneus* | F | striped | b | b | b | b | b | b | m | m |
| *subterraneus* | Q | striped | y | b | y | b | b | b | m | m |
| *subterraneus* | M | striped | y | b | y | y | b | y | y | y |
| *subterraneus* | F | white_tailed | y | b | b | b | b | b | w | w |
| *subterraneus* | Q | white_tailed | y | b | y | b | b | b | w | w |
| *sylvarum* | M | red_tailed | y | b | y | m | m | m | r | r |

| | | | | | | | | | | |
|---|---|---|---|---|---|---|---|---|---|---|
| *sylvarum* | F | red_tailed | y | b | y | m | m | m | r | r |
| *sylvarum* | Q | red_tailed | y | b | y | m | m | m | r | r |
| *sylvestris* | M | white_tailed | y | b | b | m | b | y | y | m |
| *sylvestris* | Q | white_tailed | y | b | b | m | b | m | w | b |
| *terrestris* | M | white_tailed | y | b | b | b | y | b | w | w |
| *terrestris* | F | white_tailed | y | b | b | b | y | b | w | w |
| *terrestris* | Q | white_tailed | y | b | b | b | y | b | w | w |
| *vestalis* | M | white_tailed | y | b | b | b | b | y | w | w |
| *vestalis* | Q | white_tailed | y | b | b | b | b | y | w | w |
| *veteranus* | M | striped | w | b | w | m | m | m | m | m |
| *veteranus* | F | striped | w | b | w | m | m | m | m | m |
| *veteranus* | Q | striped | w | b | w | m | m | m | m | m |
| *wurflenii* | M | red_tailed | r | b | b | b | b | b | r | r |
| *wurflenii* | F | red_tailed | b | b | b | b | b | b | y | y |
| *wurflenii* | Q | red_tailed | b | b | b | b | b | b | y | y |
| *wurflenii* | M | red_tailed | y | b | b | y | y | b | r | r |
| *wurflenii* | F | red_tailed | y | b | b | y | y | b | y | y |

*) NOTES for Appendix Table IV.

**Sex**: M = male, F = female (worker), Q = queen

**Colour abbreviations**:
- Black (b): solid black;
- White (w): pure white, not yellow or cream, includes silver and grey;
- Yellow (y): bright yellow, cream, pale yellow, off-white;
- Rufous (r): includes red, orange, brown, rust;
- Multi (m): the region in question is two or more colors, either as a mix of multi-colored hairs or discrete patches of different colors; or the region in question is highly variable between individuals of the same species.

**Meaning of coloration columns** (see also figure):
- Anterior thoracic colour: the predominant colour of the hairs on the front of the thorax (scutum, anterior to tegula).
- Central thoracic colour: the predominant colour of the hairs on the central region of the thorax (part of scutum, scutellum).
- Posterior thoracic colour: the predominant colour of the hairs on the rear of the thorax (metanotum, propodium).
- T1 colour: the predominant colour of the hairs on the first upper abdominal segment (tergite 1)
- T2 colour: the predominant colour of the hairs on the second upper abdominal segment (tergite 2)
- T3 colour: the predominant colour of the hairs on the third upper abdominal segment (tergite 3)
- T4 colour: the predominant colour of the hairs on the fourth upper abdominal segment (tergite 4)
- T5+ colour: the predominant colour of the hairs on the last upper abdominal segment(s) (tergites 5, 6, & 7)

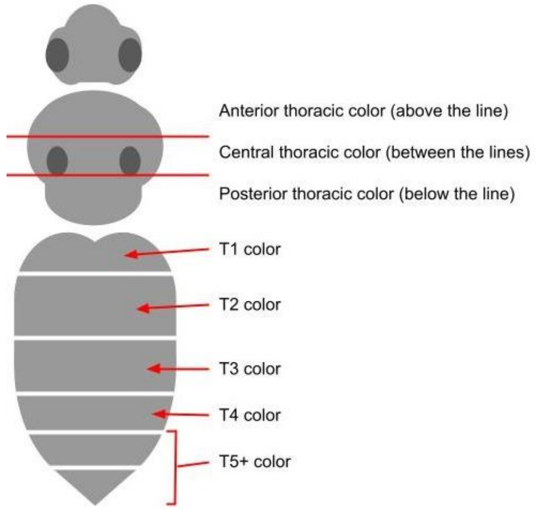

**Appendix Table V. Counts of bounding boxes containing a flower visitor in the order Diptera**

| Id. | Family | Cluster genera | Genus | Species | Nr. box | % box |
|---|---|---|---|---|---|---|
| 1 | NA | | | | 2,897 | 48.583 |
| 2 | Anthomyiidae | | *Anthomyia* | | 17 | 0.285 |
| 3 | Calliphoridae/Muscidae | | | | 184 | 3.086 |
| 4 | Chyromyidae | | | | 57 | 0.956 |
| 5 | Sarcophagidae/Tachinidae | | | | 15 | 0.252 |
| 6 | Syrphidae | | | | 7 | 0.117 |
| 7 | Syrphidae | | *Episyrphus* | *balteatus* | 224 | 3.756 |
| 8 | Syrphidae | | *Eristalis* | | 341 | 5.719 |
| 9 | Syrphidae | | *Mesembrius* | *peregrinus* | 11 | 0.184 |
| 10 | Syrphidae | | *Myathropa* | *florea* | 671 | 11.253 |
| 11 | Syrphidae | | *Pipiza* | | 4 | 0.067 |
| 12 | Syrphidae | | *Scaeva* | *pyrasti* | 4 | 0.067 |
| 13 | Syrphidae | | *Sphaerophoria* | | 289 | 4.847 |
| 14 | Syrphidae | | *Syritta* | *pipiens* | 4 | 0.067 |
| 15 | Syrphidae | black_yellow_elongated | | | 621 | 10.414 |
| 16 | Syrphidae | black_yellow_rounded | | | 488 | 8.184 |
| 17 | Syrphidae | dark_yellow_patterned | | | 38 | 0.637 |
| 18 | Syrphidae | helophilus/ parahelophilus | | | 12 | 0.201 |
| 19 | Syrphidae | small_black | | | 19 | 0.319 |
| 20 | Syrphidae | syritta/tropidia | | | 28 | 0.470 |
| 21 | Tachinidae | | *Eliozeta/Clytiomya* | | 32 | 0.537 |
| | **TOTAL** | | | | **5,963** | **100** |

**Appendix Table VI. Syrphid species grouped in clusters based on observable traits in smartphone images.**

These are species expected within Germany, listed in Ssymank et. al. 2011 at https://www.rote-liste-zentrum.de/de/Schwebfliegen-Diptera-Syrphidae-1756.html (last checked 2023-11-30).

| Genus | Species | Red list category | Custom classification name for AI |
|---|---|---|---|
| *Anasimyia* | *contracta* | threatened | Anasimyia/Lejops |
| *Anasimyia* | *interpuncta* | near_threatened | Anasimyia/Lejops |
| *Anasimyia* | *lineata* | not_threatened | Anasimyia/Lejops |
| *Anasimyia* | *lunulata* | threatened_with_extinction | Anasimyia/Lejops |
| *Anasimyia* | *transfuga* | highly_threatened | Anasimyia/Lejops |
| *Arctophila* | *bombiformis* | near_threatened | bumblebee_mimic |
| *Arctophila* | *superbiens* | threatened | Arctophila_superbiens |
| *Baccha* | *elongata* | not_threatened | Baccha |
| *Baccha* | *obscuripennis* | data_deficient | Baccha |
| *Blera* | *fallax* | not_threatened | Blera_fallax |
| *Brachyopa* | *bicolor* | threatened | Brachyopa/Hammerschmidtia |
| *Brachyopa* | *bimaculosa* | data_deficient | Brachyopa/Hammerschmidtia |
| *Brachyopa* | *dorsata* | not_threatened | Brachyopa/Hammerschmidtia |
| *Brachyopa* | *grunewaldensis* | highly_threatened | Brachyopa/Hammerschmidtia |
| *Brachyopa* | *insensilis* | not_threatened | Brachyopa/Hammerschmidtia |
| *Brachyopa* | *maculipennis* | threatened_with_extinction | Brachyopa/Hammerschmidtia |
| *Brachyopa* | *obscura* | extremely_rare | Brachyopa/Hammerschmidtia |

| | | | |
|---|---|---|---|
| *Brachyopa* | *panzeri* | not_threatened | Brachyopa/Hammerschmidtia |
| *Brachyopa* | *pilosa* | not_threatened | Brachyopa/Hammerschmidtia |
| *Brachyopa* | *plena* | data_deficient | Brachyopa/Hammerschmidtia |
| *Brachyopa* | *scutellaris* | near_threatened | Brachyopa/Hammerschmidtia |
| *Brachyopa* | *silviae* | data_deficient | Brachyopa/Hammerschmidtia |
| *Brachyopa* | *testacea* | not_threatened | Brachyopa/Hammerschmidtia |
| *Brachyopa* | *vittata* | not_threatened | Brachyopa/Hammerschmidtia |
| *Brachypalpoides* | *lentus* | not_threatened | Chalcosyrphus/Brachypalpoides |
| *Brachypalpus* | *chrysites* | threatened | honeybee_mimic |
| *Brachypalpus* | *laphriformis* | not_threatened | honeybee_mimic |
| *Brachypalpus* | *valgus* | not_threatened | honeybee_mimic |
| *Caliprobola* | *speciosa* | not_threatened | Caliprobola_speciosa |
| *Callicera* | *aenea* | threatened | Callicera |
| *Callicera* | *aurata* | threatened_with_extinction | Callicera |
| *Callicera* | *fagesii* | highly_threatened | Callicera |
| *Callicera* | *macquartii* | threatened_with_extinction | Callicera |
| *Callicera* | *rufa* | highly_threatened | Callicera |
| *Callicera* | *spinolae* | threatened_with_extinction | Callicera |
| *Ceriana* | *conopsoides* | highly_threatened | Ceriana/Sphiximorpha/Temnostoma |
| *Chalcosyrphus* | *eunotus* | highly_threatened | honeybee_mimic |
| *Chalcosyrphus* | *femoratus* | threatened | Chalcosyrphus |
| *Chalcosyrphus* | *nemorum* | not_threatened | dark_yellow_patterned |
| *Chalcosyrphus* | *piger* | highly_threatened | Chalcosyrphus/Brachypalpoides |
| *Chalcosyrphus* | *valgus* | threatened | Chalcosyrphus |
| *Chamaesyrphus* | *caledonicus* | threatened_with_extinction | dark_yellow_patterned |
| *Chamaesyrphus* | *lusitanicus* | threatened_with_extinction | dark_yellow_patterned |
| *Chamaesyrphus* | *scaevoides* | highly_threatened | dark_yellow_patterned |
| *Cheilosia* | *aerea* | Threat_of_unknown_extent | small_black |
| *Cheilosia* | *ahenea* | threatened | small_black |
| *Cheilosia* | *alba* | data_deficient | small_black |
| *Cheilosia* | *albipila* | not_threatened | small_black |
| *Cheilosia* | *albitarsis* | not_threatened | small_black |
| *Cheilosia* | *antiqua* | near_threatened | small_black |
| *Cheilosia* | *barbata* | not_threatened | small_black |
| *Cheilosia* | *bergenstammi* | not_threatened | small_black |
| *Cheilosia* | *brachysoma* | threatened_with_extinction | small_black |
| *Cheilosia* | *bracusi* | data_deficient | small_black |
| *Cheilosia* | *caerulescens* | not_threatened | small_black |
| *Cheilosia* | *canicularis* | not_threatened | small_black |
| *Cheilosia* | *carbonaria* | not_threatened | small_black |
| *Cheilosia* | *chlorus* | not_threatened | small_black |
| *Cheilosia* | *chrysocoma* | not_threatened | Cheilosia_chrysocoma |
| *Cheilosia* | *clama* | data_deficient | small_black |
| *Cheilosia* | *crassiseta* | not_threatened | small_black |
| *Cheilosia* | *cynocephala* | data_deficient | small_black |
| *Cheilosia* | *derasa* | not_threatened | small_black |
| *Cheilosia* | *fasciata* | not_threatened | small_black |
| *Cheilosia* | *faucis* | not_threatened | small_black |

| | | | |
|---|---|---|---|
| *Cheilosia* | *flavipes* | not_threatened | small_black |
| *Cheilosia* | *fraterna* | not_threatened | small_black |
| *Cheilosia* | *frontalis* | not_threatened | small_black |
| *Cheilosia* | *gagatea* | extremely_rare | small_black |
| *Cheilosia* | *gigantea* | not_threatened | small_black |
| *Cheilosia* | *griseifacies* | threatened_with_extinction | small_black |
| *Cheilosia* | *grisella* | threatened | small_black |
| *Cheilosia* | *grossa* | not_threatened | Cheilosia_grossa |
| *Cheilosia* | *hercyniae* | extremely_rare | small_black |
| *Cheilosia* | *himantopus* | not_threatened | small_black |
| *Cheilosia* | *illustrata* | not_threatened | Cheilosia_illustrata |
| *Cheilosia* | *impressa* | not_threatened | small_black |
| *Cheilosia* | *impudens* | threatened | small_black |
| *Cheilosia* | *insignis* | extremely_rare | small_black |
| *Cheilosia* | *laeviseta* | extremely_rare | small_black |
| *Cheilosia* | *laeviventris* | extremely_rare | small_black |
| *Cheilosia* | *lasiopa* | not_threatened | small_black |
| *Cheilosia* | *laticornis* | highly_threatened | small_black |
| *Cheilosia* | *latifrons* | not_threatened | small_black |
| *Cheilosia* | *lenis* | not_threatened | small_black |
| *Cheilosia* | *loewi* | threatened_with_extinction | small_black |
| *Cheilosia* | *longula* | near_threatened | small_black |
| *Cheilosia* | *melanopa* | not_threatened | small_black |
| *Cheilosia* | *melanura* | not_threatened | small_black |
| *Cheilosia* | *montana* | not_threatened | small_black |
| *Cheilosia* | *morio* | not_threatened | small_black |
| *Cheilosia* | *mutabilis* | Threat_of_unknown_extent | small_black |
| *Cheilosia* | *nebulosa* | threatened | small_black |
| *Cheilosia* | *nigripes* | not_threatened | small_black |
| *Cheilosia* | *nivalis* | not_threatened | small_black |
| *Cheilosia* | *orthotricha* | near_threatened | small_black |
| *Cheilosia* | *pagana* | not_threatened | small_black |
| *Cheilosia* | *pascuorum* | highly_threatened | small_black |
| *Cheilosia* | *pedemontana* | not_threatened | small_black |
| *Cheilosia* | *personata* | threatened | Cheilosia_personata |
| *Cheilosia* | *pictipennis* | extremely_rare | small_black |
| *Cheilosia* | *pilifer* | extremely_rare | small_black |
| *Cheilosia* | *pini* | threatened_with_extinction | small_black |
| *Cheilosia* | *proxima* | not_threatened | small_black |
| *Cheilosia* | *psilophthalma* | near_threatened | small_black |
| *Cheilosia* | *pubera* | threatened | small_black |
| *Cheilosia* | *ranunculi* | near_threatened | small_black |
| *Cheilosia* | *rhynchops* | not_threatened | small_black |
| *Cheilosia* | *ruficollis* | NA | small_black |
| *Cheilosia* | *rufimana* | threatened | small_black |
| *Cheilosia* | *sahlbergi* | not_threatened | small_black |
| *Cheilosia* | *scanica* | not_threatened | small_black |
| *Cheilosia* | *scutellata* | not_threatened | small_black |

| | | | |
|---|---|---|---|
| *Cheilosia* | *semifasciata* | not_threatened | small_black |
| *Cheilosia* | *soror* | not_threatened | small_black |
| *Cheilosia* | *subpictipennis* | threatened | small_black |
| *Cheilosia* | *urbana* | not_threatened | small_black |
| *Cheilosia* | *uviformis* | data_deficient | small_black |
| *Cheilosia* | *variabilis* | not_threatened | small_black |
| *Cheilosia* | *velutina* | not_threatened | small_black |
| *Cheilosia* | *venosa* | extremely_rare | small_black |
| *Cheilosia* | *vernalis* | not_threatened | small_black |
| *Cheilosia* | *vicina* | not_threatened | small_black |
| *Cheilosia* | *vulpina* | not_threatened | small_black |
| *Chrysogaster* | *basalis* | threatened | Chrysogaster |
| *Chrysogaster* | *cemiteriorum* | threatened | small_black |
| *Chrysogaster* | *rondanii* | Threat_of_unknown_extent | small_black |
| *Chrysogaster* | *solstitialis* | not_threatened | Chrysogaster |
| *Chrysogaster* | *virescens* | Threat_of_unknown_extent | small_black |
| *Chrysotoxum* | *arcuatum* | not_threatened | wasp_mimic_seperated_stripes |
| *Chrysotoxum* | *bicinctum* | not_threatened | Chrysotoxum_bicinctum |
| *Chrysotoxum* | *cautum* | not_threatened | wasp_mimic_seperated_stripes |
| *Chrysotoxum* | *elegans* | threatened_with_extinction | wasp_mimic_seperated_stripes |
| *Chrysotoxum* | *fasciolatum* | near_threatened | wasp_mimic_seperated_stripes |
| *Chrysotoxum* | *festivum* | not_threatened | wasp_mimic_seperated_stripes |
| *Chrysotoxum* | *intermedium* | not_threatened | wasp_mimic_seperated_stripes |
| *Chrysotoxum* | *lineare* | threatened_with_extinction | wasp_mimic_seperated_stripes |
| *Chrysotoxum* | *octomaculatum* | threatened_with_extinction | wasp_mimic_seperated_stripes |
| *Chrysotoxum* | *vernale* | not_threatened | wasp_mimic_seperated_stripes |
| *Chrysotoxum* | *verralli* | not_threatened | wasp_mimic_seperated_stripes |
| *Criorhina* | *asilica* | not_threatened | honeybee_mimic |
| *Criorhina* | *pachymera* | threatened_with_extinction | honeybee_mimic |
| *Criorhina* | *ranunculi* | not_threatened | bumblebee_mimic |
| *Criorhina* | *berberina* | not_threatened | bumblebee_mimic |
| *Criorhina* | *floccosa* | Threat_of_unknown_extent | bumblebee_mimic |
| *Dasysyrphus* | *albostriatus* | not_threatened | black_yellow_rounded |
| *Dasysyrphus* | *friuliensis* | not_threatened | black_yellow_rounded |
| *Dasysyrphus* | *hilaris* | not_threatened | black_yellow_rounded |
| *Dasysyrphus* | *lenensis* | not_threatened | black_yellow_rounded |
| *Dasysyrphus* | *pauxillus* | not_threatened | black_yellow_rounded |
| *Dasysyrphus* | *pinastri* | not_threatened | black_yellow_rounded |
| *Dasysyrphus* | *postclaviger* | data_deficient | black_yellow_rounded |
| *Dasysyrphus* | *tricinctus* | not_threatened | Dasysyrphus_tricinctus |
| *Dasysyrphus* | *venustus* | not_threatened | black_yellow_rounded |
| *Didea* | *alneti* | near_threatened | black_yellow_rounded |
| *Didea* | *fasciata* | not_threatened | black_yellow_rounded |
| *Didea* | *intermedia* | not_threatened | black_yellow_rounded |
| *Doros* | *profuges* | highly_threatened | Doros_profuges |
| *Epistrophe* | *cryptica* | not_threatened | black_yellow_rounded |
| *Epistrophe* | *diaphana* | not_threatened | black_yellow_rounded |
| *Epistrophe* | *eligans* | not_threatened | black_yellow_rounded |

| | | | |
|---|---|---|---|
| *Epistrophe* | *flava* | not_threatened | black_yellow_rounded |
| *Epistrophe* | *grossulariae* | not_threatened | black_yellow_rounded |
| *Epistrophe* | *leiophthalma* | extremely_rare | black_yellow_rounded |
| *Epistrophe* | *melanostoma* | not_threatened | black_yellow_rounded |
| *Epistrophe* | *nitidicollis* | not_threatened | black_yellow_rounded |
| *Epistrophe* | *obscuripes* | data_deficient | black_yellow_rounded |
| *Epistrophe* | *ochrostoma* | data_deficient | black_yellow_rounded |
| *Epistrophe* | *olgae* | data_deficient | black_yellow_rounded |
| *Epistrophella* | *euchroma* | not_threatened | black_yellow_elongated |
| *Episyrphus* | *balteatus* | not_threatened | Episyrphus_balteatus |
| *Eriozona* | *syrphoides* | not_threatened | bumblebee_mimic |
| *Eristalinus* | *aeneus* | not_threatened | Eristalinus |
| *Eristalinus* | *sepulchralis* | not_threatened | Eristalinus |
| *Eristalis* | *abusiva* | Threat_of_unknown_extent | Eristalis |
| *Eristalis* | *alpina* | threatened | Eristalis |
| *Eristalis* | *anthophorina* | threatened_with_extinction | bumblebee_mimic |
| *Eristalis* | *arbustorum* | not_threatened | Eristalis |
| *Eristalis* | *cryptarum* | threatened_with_extinction | Eristalis cryptarum |
| *Eristalis* | *horticola* | not_threatened | Eristalis_horticola |
| *Eristalis* | *intricaria* | not_threatened | bumblebee_mimic |
| *Eristalis* | *jugorum* | not_threatened | Eristalis |
| *Eristalis* | *nemorum* | not_threatened | Eristalis |
| *Eristalis* | *oestracea* | threatened_with_extinction | Eristalis oestracea |
| *Eristalis* | *pertinax* | not_threatened | Eristalis |
| *Eristalis* | *picea* | not_threatened | Eristalis |
| *Eristalis* | *pseudorupium* | highly_threatened | Eristalis |
| *Eristalis* | *rupium* | not_threatened | Eristalis |
| *Eristalis* | *similis* | not_threatened | Eristalis |
| *Eristalis* | *tenax* | not_threatened | Eristalis |
| *Eumerus* | *amoenus* | threatened_with_extinction | small_black |
| *Eumerus* | *clavatus* | threatened_with_extinction | small_black |
| *Eumerus* | *flavitarsis* | not_threatened | small_black |
| *Eumerus* | *grandis* | extinct | NA |
| *Eumerus* | *longicornis* | threatened_with_extinction | small_black |
| *Eumerus* | *ornatus* | not_threatened | small_black |
| *Eumerus* | *ovatus* | threatened_with_extinction | small_black |
| *Eumerus* | *ruficornis* | threatened_with_extinction | small_black |
| *Eumerus* | *sabulonum* | highly_threatened | small_black |
| *Eumerus* | *sinuatus* | threatened_with_extinction | small_black |
| *Eumerus* | *sogdianus* | data_deficient | small_black |
| *Eumerus* | *strigatus* | not_threatened | small_black |
| *Eumerus* | *tarsalis* | highly_threatened | small_black |
| *Eumerus* | *tricolor* | threatened | small_black |
| *Eumerus* | *tuberculatus* | not_threatened | small_black |
| *Eumerus* | *uncipes* | extinct | NA |
| *Eupeodes* | *bucculatus* | not_threatened | black_yellow_rounded |
| *Eupeodes* | *corollae* | not_threatened | black_yellow_rounded |
| *Eupeodes* | *goeldlini* | data_deficient | black_yellow_rounded |

| | | | |
|---|---|---|---|
| *Eupeodes* | *latifasciatus* | not_threatened | black_yellow_rounded |
| *Eupeodes* | *lundbecki* | Threat_of_unknown_extent | black_yellow_rounded |
| *Eupeodes* | *luniger* | not_threatened | black_yellow_rounded |
| *Eupeodes* | *nielseni* | not_threatened | black_yellow_rounded |
| *Eupeodes* | *nitens* | not_threatened | black_yellow_rounded |
| *Melangyna* | *cinctus* | not_threatened | black_yellow_elongated |
| *Ferdinandea* | *cuprea* | not_threatened | Ferdinandea_cuprea |
| *Ferdinandea* | *ruficornis* | threatened | Ferdinandea_ruficornis |
| *Hammerschmidtia* | *ferruginea* | threatened_with_extinction | Brachyopa/Hammerschmidtia |
| *Helophilus* | *affinis* | not_threatened | Helophilus/Parahelophilus |
| *Helophilus* | *hybridus* | not_threatened | Helophilus/Parahelophilus |
| *Helophilus* | *pendulus* | not_threatened | Helophilus/Parahelophilus |
| *Helophilus* | *trivittatus* | not_threatened | Helophilus/Parahelophilus |
| *Heringia* | *brevidens* | Threat_of_unknown_extent | small_black |
| *Heringia* | *heringi* | not_threatened | small_black |
| *Heringia* | *larusi* | data_deficient | small_black |
| *Heringia* | *latitarsis* | not_threatened | small_black |
| *Heringia* | *pubescens* | not_threatened | small_black |
| *Heringia* | *senilis* | data_deficient | small_black |
| *Heringia* | *verrucula* | extremely_rare | small_black |
| *Heringia* | *vitripennis* | not_threatened | small_black |
| *Eupeodes* | *lapponicus* | not_threatened | Eupeodes |
| *Lejogaster* | *metallina* | near_threatened | small_black |
| *Lejogaster* | *tarsata* | highly_threatened | small_black |
| *Lejops* | *vittatus* | threatened_with_extinction | Anasimyia_Lejops |
| *Lejota* | *ruficornis* | highly_threatened | Anasimyia_Lejops |
| *Leucozona* | *glaucia* | near_threatened | Leucozona_glaucia |
| *Leucozona* | *laternaria* | near_threatened | Leucozona_laternaria |
| *Leucozona* | *inopinata* | not_threatened | Leucozona |
| *Leucozona* | *lucorum* | not_threatened | Leucozona |
| *Mallota* | *cimbiciformis* | highly_threatened | honeybee_mimic |
| *Mallota* | *fuciformis* | threatened | bumblebee_mimic |
| *Mallota* | *megilliformis* | extinct | NA |
| *Megasyrphus* | *erraticus* | not_threatened | black_yellow_rounded |
| *Melangyna* | *arctica* | extremely_rare | black_yellow_elongated |
| *Melangyna* | *barbifrons* | not_threatened | black_yellow_elongated |
| *Melangyna* | *compositarum* | not_threatened | black_yellow_elongated |
| *Melangyna* | *ericarum* | threatened_with_extinction | black_yellow_elongated |
| *Melangyna* | *labiatarum* | not_threatened | black_yellow_elongated |
| *Melangyna* | *lasiophthalma* | not_threatened | black_yellow_elongated |
| *Melangyna* | *lucifera* | not_threatened | black_yellow_elongated |
| *Melangyna* | *quadrimaculata* | not_threatened | black_yellow_elongated |
| *Melangyna* | *umbellatarum* | not_threatened | black_yellow_elongated |
| *Melanogaster* | *aerosa* | highly_threatened | small_black |
| *Melanogaster* | *curvistylus* | threatened_with_extinction | small_black |
| *Melanogaster* | *hirtella* | not_threatened | small_black |
| *Melanogaster* | *nuda* | not_threatened | small_black |
| *Melanogaster* | *parumplicata* | highly_threatened | small_black |

| | | | |
|---|---|---|---|
| *Melanostoma* | *alpinum* | NA | dark_yellow_patterned |
| *Melanostoma* | *dubium* | threatened | dark_yellow_patterned |
| *Melanostoma* | *mellinum* | not_threatened | dark_yellow_patterned |
| *Melanostoma* | *scalare* | not_threatened | dark_yellow_patterned |
| *Meligramma* | *cingulata* | not_threatened | black_yellow_elongated |
| *Meligramma* | *guttatum* | Threat_of_unknown_extent | black_yellow_elongated |
| *Meligramma* | *trianguliferum* | not_threatened | black_yellow_elongated |
| *Meliscaeva* | *auricollis* | not_threatened | black_yellow_elongated |
| *Meliscaeva* | *cinctella* | not_threatened | black_yellow_elongated |
| *Merodon* | *aberrans* | threatened_with_extinction | honeybee_mimic |
| *Merodon* | *aeneus* | threatened_with_extinction | honeybee_mimic |
| *Merodon* | *armipes* | highly_threatened | honeybee_mimic |
| *Merodon* | *avidus* | near_threatened | Merodon |
| *Merodon* | *cinereus* | threatened | Merodon |
| *Merodon* | *constans* | threatened_with_extinction | honeybee_mimic |
| *Merodon* | *equestris* | not_threatened | bumblebee_mimic |
| *Merodon* | *nigritarsis* | extremely_rare | Merodon |
| *Merodon* | *ruficornis* | highly_threatened | Merodon |
| *Merodon* | *rufus* | near_threatened | honeybee_mimic |
| *Mesembrius* | *peregrinus* | threatened_with_extinction | Mesembrius peregrinus |
| *Microdon* | *analis* | not_threatened | Microdon |
| *Microdon* | *devius* | near_threatened | Microdon |
| *Microdon* | *major* | data_deficient | Microdon |
| *Microdon* | *miki* | threatened_with_extinction | Microdon |
| *Microdon* | *mutabilis* | data_deficient | Microdon |
| *Microdon* | *myrmicae* | data_deficient | Microdon |
| *Myathropa* | *florea* | not_threatened | Myathropa_florea |
| *Myolepta* | *dubia* | near_threatened | Myolepta |
| *Myolepta* | *obscura* | threatened_with_extinction | Myolepta |
| *Myolepta* | *potens* | highly_threatened | Myolepta |
| *Myolepta* | *vara* | threatened | Myolepta |
| *Neoascia* | *annexa* | not_threatened | Neoascia/Sphegina/Spazigaster |
| *Neoascia* | *geniculata* | highly_threatened | Neoascia/Sphegina/Spazigaster |
| *Neoascia* | *interrupta* | near_threatened | Neoascia/Sphegina/Spazigaster |
| *Neoascia* | *meticulosa* | not_threatened | Neoascia/Sphegina/Spazigaster |
| *Neoascia* | *obliqua* | not_threatened | Neoascia/Sphegina/Spazigaster |
| *Neoascia* | *podagrica* | not_threatened | Neoascia/Sphegina/Spazigaster |
| *Neoascia* | *tenur* | not_threatened | Neoascia/Sphegina/Spazigaster |
| *Neoascia* | *unifasciata* | near_threatened | Neoascia/Sphegina/Spazigaster |
| *Orthonevra* | *brevicornis* | near_threatened | small_black |
| *Orthonevra* | *elegans* | threatened_with_extinction | small_black |
| *Orthonevra* | *erythrogona* | threatened_with_extinction | small_black |
| *Orthonevra* | *frontalis* | extinct | NA |
| *Orthonevra* | *geniculata* | threatened | small_black |
| *Orthonevra* | *incisa* | threatened_with_extinction | small_black |
| *Orthonevra* | *intermedia* | threatened | Orthonevra_intermedia |
| *Orthonevra* | *montana* | extremely_rare | small_black |
| *Orthonevra* | *nobilis* | not_threatened | small_black |

| | | | |
|---|---|---|---|
| *Orthonevra* | *stackelbergi* | threatened_with_extinction | small_black |
| *Orthonevra* | *tristis* | Threat_of_unknown_extent | small_black |
| *Paragus* | *albifrons* | threatened | small_black |
| *Paragus* | *bicolor* | near_threatened | small_black |
| *Paragus* | *constrictus* | data_deficient | small_black |
| *Paragus* | *finitimus* | Threat_of_unknown_extent | Paragus |
| *Paragus* | *flammeus* | highly_threatened | small_black |
| *Paragus* | *haemorrhous* | not_threatened | Paragus |
| *Paragus* | *majoranae* | threatened_with_extinction | small_black |
| *Paragus* | *pecchiolii* | not_threatened | small_black |
| *Paragus* | *punctulatus* | not_threatened | small_black |
| *Paragus* | *quadrifasciatus* | not_threatened | Paragus_quadrifasciatus |
| *Paragus* | *tibialis* | highly_threatened | small_black |
| *Parasyrphus* | *annulatus* | not_threatened | black_yellow_rounded |
| *Parasyrphus* | *lineola* | not_threatened | black_yellow_rounded |
| *Parasyrphus* | *macularis* | not_threatened | black_yellow_rounded |
| *Parasyrphus* | *malinellus* | not_threatened | black_yellow_rounded |
| *Parasyrphus* | *nigritarsis* | data_deficient | black_yellow_rounded |
| *Parasyrphus* | *punctulatus* | not_threatened | black_yellow_rounded |
| *Parasyrphus* | *vittiger* | not_threatened | black_yellow_rounded |
| *Parhelophilus* | *consimilis* | highly_threatened | Helophilus/Parahelophilus |
| *Parhelophilus* | *frutetorum* | near_threatened | Helophilus/Parahelophilus |
| *Parhelophilus* | *versicolor* | near_threatened | Helophilus/Parahelophilus |
| *Pelecocera* | *tricincta* | threatened | dark_yellow_patterned |
| *Pipiza* | *accola* | threatened_with_extinction | Pipiza |
| *Pipiza* | *austriaca* | not_threatened | Pipiza |
| *Pipiza* | *bimaculata* | not_threatened | Pipiza |
| *Pipiza* | *fenestrata* | data_deficient | Pipiza |
| *Pipiza* | *festiva* | near_threatened | Pipiza |
| *Pipiza* | *lugubris* | not_threatened | Pipiza |
| *Pipiza* | *luteitarsis* | threatened | Pipiza |
| *Pipiza* | *noctiluca* | not_threatened | Pipiza |
| *Pipiza* | *notata* | NA | Pipiza |
| *Pipiza* | *quadrimaculata* | not_threatened | Pipiza |
| *Pipiza* | *signata* | NA | Pipiza |
| *Pipizella* | *annulata* | near_threatened | small_black |
| *Pipizella* | *divicoi* | not_threatened | small_black |
| *Pipizella* | *maculipennis* | extinct | NA |
| *Pipizella* | *mongolorum* | threatened_with_extinction | small_black |
| *Pipizella* | *nigriana* | not_threatened | small_black |
| *Pipizella* | *pennina* | threatened_with_extinction | small_black |
| *Pipizella* | *viduata* | not_threatened | small_black |
| *Pipizella* | *virens* | Threat_of_unknown_extent | small_black |
| *Pipizella* | *zeneggenensis* | near_threatened | small_black |
| *Platycheirus* | *abruzzensis* | extremely_rare | dark_yellow_patterned |
| *Platycheirus* | *albimanus* | not_threatened | dark_yellow_patterned |
| *Platycheirus* | *ambiguus* | Threat_of_unknown_extent | dark_yellow_patterned |
| *Platycheirus* | *amplus* | threatened_with_extinction | dark_yellow_patterned |

| | | | |
|---|---|---|---|
| *Platycheirus* | *angustatus* | not_threatened | dark_yellow_patterned |
| *Platycheirus* | *angustipes* | Threat_of_unknown_extent | dark_yellow_patterned |
| *Platycheirus* | *aurolateralis* | data_deficient | dark_yellow_patterned |
| *Platycheirus* | *clypeatus* | not_threatened | dark_yellow_patterned |
| *Platycheirus* | *complicatus* | not_threatened | dark_yellow_patterned |
| *Platycheirus* | *discimanus* | not_threatened | dark_yellow_patterned |
| *Platycheirus* | *europaeus* | not_threatened | dark_yellow_patterned |
| *Platycheirus* | *fasciculatus* | extremely_rare | dark_yellow_patterned |
| *Platycheirus* | *fulviventris* | near_threatened | dark_yellow_patterned |
| *Platycheirus* | *immaculatus* | not_threatened | dark_yellow_patterned |
| *Platycheirus* | *immarginatus* | threatened_with_extinction | dark_yellow_patterned |
| *Platycheirus* | *jaerensis* | extremely_rare | dark_yellow_patterned |
| *Platycheirus* | *laskai* | extremely_rare | dark_yellow_patterned |
| *Platycheirus* | *manicatus* | not_threatened | dark_yellow_patterned |
| *Platycheirus* | *melanopsis* | not_threatened | dark_yellow_patterned |
| *Platycheirus* | *nielseni* | not_threatened | dark_yellow_patterned |
| *Platycheirus* | *occultus* | near_threatened | dark_yellow_patterned |
| *Platycheirus* | *parmatus* | not_threatened | dark_yellow_patterned |
| *Platycheirus* | *peltatus* | not_threatened | dark_yellow_patterned |
| *Platycheirus* | *perpallidus* | threatened | dark_yellow_patterned |
| *Platycheirus* | *podagratus* | highly_threatened | dark_yellow_patterned |
| *Platycheirus* | *scambus* | near_threatened | dark_yellow_patterned |
| *Platycheirus* | *scutatus* | not_threatened | dark_yellow_patterned |
| *Platycheirus* | *splendidus* | data_deficient | dark_yellow_patterned |
| *Platycheirus* | *sticticus* | Threat_of_unknown_extent | dark_yellow_patterned |
| *Platycheirus* | *tarsalis* | not_threatened | dark_yellow_patterned |
| *Platycheirus* | *tatricus* | not_threatened | dark_yellow_patterned |
| *Platycheirus* | *transfugus* | threatened_with_extinction | dark_yellow_patterned |
| *Pocota* | *personata* | threatened_with_extinction | bumblebee_mimic |
| *Portevinia* | *maculata* | not_threatened | small_black |
| *Psarus* | *abdominalis* | threatened_with_extinction | Psarus_abdominalis |
| *Psilota* | *anthracina* | data_deficient | small_black |
| *Psilota* | *atra* | data_deficient | small_black |
| *Psilota* | *innupta* | extremely_rare | small_black |
| *Pyrophaena* | *granditarsa* | near_threatened | dark_yellow_patterned |
| *Pyrophaena* | *rosarum* | not_threatened | dark_yellow_patterned |
| *Rhingia* | *borealis* | not_threatened | Rhingia |
| *Rhingia* | *campestris* | not_threatened | Rhingia |
| *Rhingia* | *rostrata* | highly_threatened | Rhingia |
| *Riponnensia* | *splendens* | highly_threatened | small_black |
| *Scaeva* | *dignota* | not_threatened | Scaeva |
| *Scaeva* | *pyrastri* | not_threatened | Scaeva |
| *Scaeva* | *selenitica* | not_threatened | Scaeva |
| *Sericomyia* | *lappona* | threatened | Sericomyia |
| *Sericomyia* | *silentis* | not_threatened | Sericomyia |
| *Spazigaster* | *ambulans* | Threat_of_unknown_extent | Neoascia/Sphegina/Spazigaster |
| *Sphaerophoria* | *bankowskae* | not_threatened | black_yellow_elongated |
| *Sphaerophoria* | *batava* | not_threatened | black_yellow_elongated |

| | | | |
|---|---|---|---|
| *Sphaerophoria* | *chongjini* | threatened | black_yellow_elongated |
| *Sphaerophoria* | *estebani* | extremely_rare | black_yellow_elongated |
| *Sphaerophoria* | *fatarum* | near_threatened | black_yellow_elongated |
| *Sphaerophoria* | *infuscata* | threatened | black_yellow_elongated |
| *Sphaerophoria* | *interrupta* | not_threatened | black_yellow_elongated |
| *Sphaerophoria* | *loewi* | threatened_with_extinction | black_yellow_elongated |
| *Sphaerophoria* | *philanthus* | threatened | black_yellow_elongated |
| *Sphaerophoria* | *potentillae* | threatened_with_extinction | black_yellow_elongated |
| *Sphaerophoria* | *rueppellii* | not_threatened | black_yellow_elongated |
| *Sphaerophoria* | *scripta* | not_threatened | black_yellow_elongated |
| *Sphaerophoria* | *shirchan* | data_deficient | black_yellow_elongated |
| *Sphaerophoria* | *taeniata* | not_threatened | black_yellow_elongated |
| *Sphaerophoria* | *virgata* | not_threatened | black_yellow_elongated |
| *Sphegina* | *clavata* | not_threatened | Neoascia/Sphegina/Spazigaster |
| *Sphegina* | *clunipes* | not_threatened | Neoascia/Sphegina/Spazigaster |
| *Sphegina* | *cornifera* | not_threatened | Neoascia/Sphegina/Spazigaster |
| *Sphegina* | *elegans* | not_threatened | Neoascia/Sphegina/Spazigaster |
| *Sphegina* | *latifrons* | not_threatened | Neoascia/Sphegina/Spazigaster |
| *Sphegina* | *montana* | not_threatened | Neoascia/Sphegina/Spazigaster |
| *Sphegina* | *platychira* | highly_threatened | Neoascia/Sphegina/Spazigaster |
| *Sphegina* | *sibirica* | not_threatened | Neoascia/Sphegina/Spazigaster |
| *Sphegina* | *spheginea* | highly_threatened | Neoascia/Sphegina/Spazigaster |
| *Sphegina* | *verecunda* | not_threatened | Neoascia/Sphegina/Spazigaster |
| *Sphiximorpha* | *binominata* | extremely_rare | Ceriana/Sphiximorpha/Temnostoma |
| *Sphiximorpha* | *subsessilis* | highly_threatened | Ceriana/Sphiximorpha/Temnostoma |
| *Spilomyia* | *digitata* | data_deficient | wasp_mimic_banded |
| *Spilomyia* | *diophthalma* | threatened_with_extinction | wasp_mimic_banded |
| *Spilomyia* | *manicata* | threatened_with_extinction | wasp_mimic_banded |
| *Syritta* | *pipiens* | not_threatened | Syritta/Tropidia |
| *Syrphocheilosia* | *claviventris* | not_threatened | small_black |
| *Syrphus* | *auberti* | extremely_rare | black_yellow_rounded |
| *Syrphus* | *nitidifrons* | not_threatened | black_yellow_rounded |
| *Syrphus* | *ribesii* | not_threatened | black_yellow_rounded |
| *Syrphus* | *torvus* | not_threatened | black_yellow_rounded |
| *Syrphus* | *vitripennis* | not_threatened | black_yellow_rounded |
| *Temnostoma* | *apiforme* | threatened | wasp_mimic_banded |
| *Temnostoma* | *bombylans* | not_threatened | Ceriana/Sphiximorpha/Temnostoma |
| *Temnostoma* | *meridionale* | threatened | Temnostoma |
| *Temnostoma* | *vespiforme* | not_threatened | Temnostoma |
| *Trichopsomyia* | *flavitarsis* | not_threatened | small_black |
| *Trichopsomyia* | *joratensis* | not_threatened | small_black |
| *Trichopsomyia* | *lucida* | threatened | small_black |
| *Triglyphus* | *primus* | not_threatened | small_black |
| *Tropidia* | *fasciata* | threatened_with_extinction | Syritta/Tropidia |
| *Tropidia* | *scita* | not_threatened | Syritta/Tropidia |
| *Volucella* | *bombylans* | not_threatened | Volucella_bombylans |
| *Volucella* | *inanis* | not_threatened | Volucella_inanis |
| *Volucella* | *inflata* | threatened | Volucella_inflata |

| | | | |
|---|---|---|---|
| *Volucella* | *pellucens* | not_threatened | Volucella_pellucens |
| *Volucella* | *zonaria* | not_threatened | Volucella_zonaria |
| *Xanthandrus* | *comtus* | not_threatened | dark_yellow_patterned |
| *Xanthogramma* | *citrofasciatum* | near_threatened | wasp_mimic_seperated_stripes |
| *Xanthogramma* | *dives* | data_deficient | wasp_mimic_seperated_stripes |
| *Xanthogramma* | *laetum* | not_threatened | wasp_mimic_seperated_stripes |
| *Xanthogramma* | *pedissequum* | not_threatened | wasp_mimic_seperated_stripes |
| *Xanthogramma* | *stackelbergi* | data_deficient | wasp_mimic_seperated_stripes |
| *Xylota* | *abiens* | not_threatened | dark_yellow_patterned |
| *Xylota* | *caeruleiventris* | threatened_with_extinction | dark_yellow_patterned |
| *Xylota* | *florum* | not_threatened | dark_yellow_patterned |
| *Xylota* | *ignava* | threatened | dark_yellow_patterned |
| *Xylota* | *jakutorum* | not_threatened | dark_yellow_patterned |
| *Xylota* | *meigeniana* | highly_threatened | dark_yellow_patterned |
| *Xylota* | *segnis* | not_threatened | dark_yellow_patterned |
| *Xylota* | *sylvarum* | not_threatened | Xylota |
| *Xylota* | *tarda* | not_threatened | dark_yellow_patterned |
| *Xylota* | *xanthocnema* | threatened | Xylota |

**Appendix Table VII. Fieldwork protocol**

This tutorial will guide you through the optimal process of setting the smartphone on top of a target flower. Specifically, this involves setting the optimal distance from the flower and minimising background noise to capture clear images of pollinators.

### A. Pre-fieldwork Checklist

Below is a list of the equipment you will need to bring to the field:

1. Tripods suitable for phones - both small and large
2. Sticks to stabilise the plants (bring a sufficient quantity)
3. Yarn to attach the plant to the stick
4. Scissors for cutting the yarn as necessary
5. Power banks for charging
6. USB cables for phone-to-power bank connection, ensure they are long and compatible with your phone models
7. Mobile phones and any associated accessories
8. Adequate quantities of printed field note forms (refer to section D.5) along with pencils
9. Chairs for rest and work
10. Adequate food and water supplies
11. Recommended: sunscreen, sunhat and rain gear (check weather forecast)
12. Check weather forecasts: umbrellas and waterproof clothing may be needed
13. Install the Flora Incognita app on your phone for plant identification. Note that this app requires an internet connection to function.

Ensure that the following devices are fully charged before fieldwork begins:

1. Mobile phones
2. Power banks. Keep an eye out for any signs of failure, such as bulging due to overheating in the sun. If this occurs, cease usage immediately and arrange for proper disposal and replacement.

Verify that there is sufficient storage space on the phones to store images. At the end of each day, download the day's images for storage and clear the phone's memory.

If you are driving, remember to bring your valid driver's licence and identification card/passport.

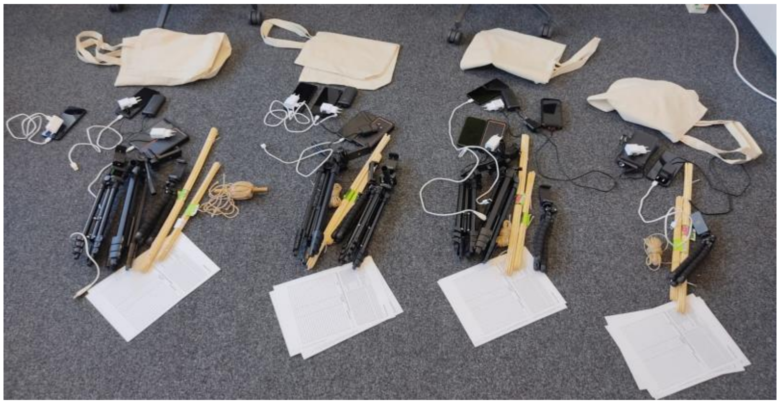

Figure 1. Example of gear for fieldwork

### B. During Fieldwork

#### B.1. Fieldwork procedure

1. Select a target flower, preferably choosing an open flower whenever possible (refer to section B.2. Selection of Open Flowers).
2. Turn on the phone if it is not already on. Confirm that the app settings are configured correctly (refer to section C.1. Settings). These should have been set at the start of the field season and should remain unchanged unless otherwise instructed. However, it is good practice to double-check that everything is correctly set.
3. For each individual flower or plant, create a separate folder on your phone (see section C.2. How to create a "plant-folder", e.g., centaurea-scabiosa-vs-01).
4. Assess the position of the sun and anticipate its path. It is crucial to avoid having the sun directly in front or behind the camera as it could lead to underexposed or overexposed images. Thus, the phone should not directly face the sun; the sun should be to your left or right. Select a flower that allows for these conditions.
5. Secure the stick firmly near the flower to prevent wind displacement. Refer to section B.3. The support stick, for more details on stick placement and usage.
6. Secure the flower to the stick without damaging the plant (refer to section B.5. Using yarn to secure the flower to the stick). Make sure the yarn is not visible in the image and is never situated between the flower and the camera, as it obstructs visibility.
7. Mount the phone onto the tripod.
8. Connect the phone to the power bank. Ensure neither the power bank nor the cable obscures the camera's view.
9. After determining the optimal position relative to the sun and the flower, ascertain the best distance from the flower (see sections B.4. Image background and distance from the flower & B.2. Selection of open flowers).
10. Confirm the stability of the tripod and the phone.
11. Ensure the camera focus is on the flower, not on the stick or background. This is vital for image quality and deserves ample time to perfect. Reconfirm that the focus is locked and not set to auto (refer to section C.1. Settings). Make sure there's no flash as well.
12. Address any other issues that could affect image quality, such as obstructions in front of the camera, smudges on the camera lens, or lens condensation.
13. You may now begin the one-hour session. Activate the Open Camera app's shutter button to start capturing time-lapse images. Set an alarm on your personal phone to remind you to stop recording after one hour. Note that the Open Camera app does not have a built-in timer (not at the moment).
14. Complete the field notes for the flower and the session, noting down the exact folder name used for the plant (e.g., centaurea-scabiosa-vs-01).
15. Start setting up another phone on a different plant.
16. Maintain distance from the target plant to avoid scaring away potential flower visitors.
17. Periodically check if the wind has displaced the flower or tripod, verify the app is still capturing images, and troubleshoot any unexpected issues (e.g., the phone shutting down due to overheating or low battery).
18. After one hour, stop the Open Camera app by pressing the shutter button . The screen might be off to conserve battery, so you might need to activate it first.

#### B.2. Selection of open flowers

Select plants with open flowers that are easily accessible for camera positioning. The phone's surface should align parallel with the flower's surface, ensuring an optimal view of the pollinator when it settles on the flower (refer to the examples below).

Preferably avoid tall plants, especially those exceeding 1 metre in height, as the wind may easily displace them even when secured to a support stick. If the target plant species tends to be tall, ensure that the support stick is firmly anchored in the soil.

Examples of desirable open flowers and high-quality images:

<table>
<tr><td>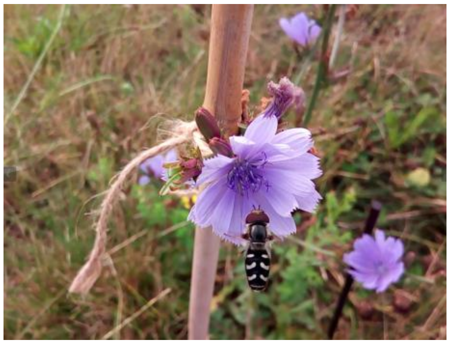<br>Image quality is good, but the shot could be closer to the flower. Note also that the support stick should be situated under the flower. A thinner stick would be preferable.</td><td>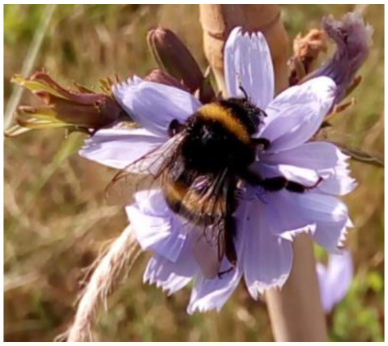</td></tr>
<tr><td>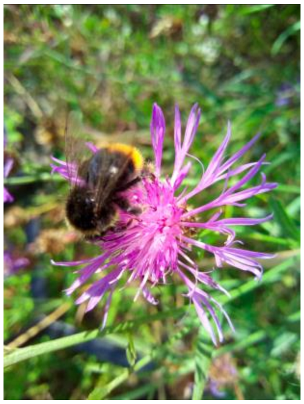<br>Both the flower and insect are in focus, occupying a significant portion of the image, allowing detailed observation of the insect's wing venation. Larger insects, like bumblebees, generally lend themselves to near-ideal images.</td><td>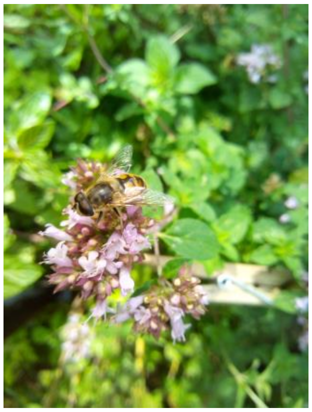<br>Good image. In an ideal scenario, there would be no artificial objects, such as the clothespin, visible in the background.</td></tr>
<tr><td>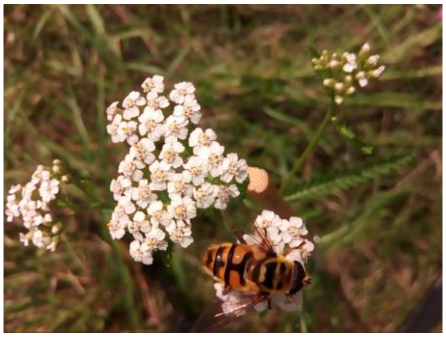<br>When dealing with inflorescences, aim to select those that aren't too wide, ensuring they can fit entirely within the photo. This allows you to capture the complete inflorescence while maintaining an optimal distance.</td><td></td></tr>
</table>

#### B.3. The support stick

Ensure that the support stick is secured firmly into the ground, ideally with its top positioned below the flower. If the top of the support stick is situated near or slightly above the flower, insects may land on it rather than on the flower. Additionally, ants may readily climb the stick, entering the camera's field of view, which we aim to minimise as much as possible.

It's important to steer clear of situations similar to those illustrated in the images below:

<table>
<tr>
<td>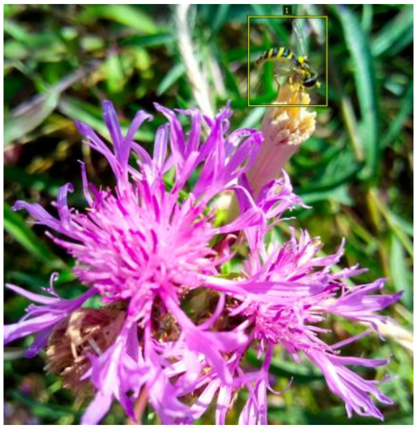<br>Insects Using the Stick for Rest: The stick should not serve as a perch for insects to rest upon.</td>
<td>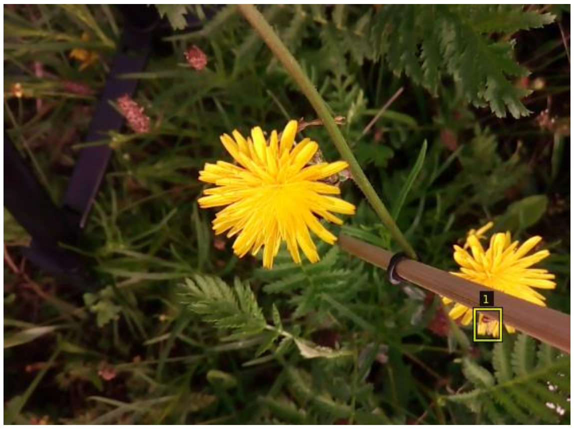<br>Stick Obscuring the Primary Flowers: The stick must not obstruct the view of the main flowers, as it limits our visibility of the insects.</td>
</tr>
<tr>
<td></td>
<td>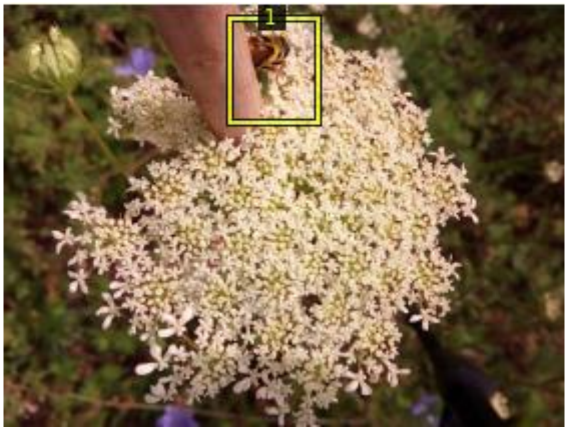<br>Insect Concealed Behind the Stick: Circumstances where the insect ends up behind the stick should be avoided. The stick should be placed directly below the flower for optimal visibility.</td>
</tr>
</table>

#### B.4. Image background and distance from the flower

Invest sufficient time in determining the optimal distance from the flower to minimise the amount of background noise. Make an effort to exclude other flowers from the background to prevent out-of-focus insects from being visible.
The flower should occupy a significant portion of the camera’s display. The objective is to make the insects appear as large as possible in the image (see examples below and refer to section "B.2. Selection of open flowers" for further examples).

Below are cases where the capture could be improved by placing the phone closer to the flower and taking photos from an overhead, rather than lateral, perspective.

<table>
<tr><th>Current situation</th><th>Possible improvements</th></tr>
<tr><td>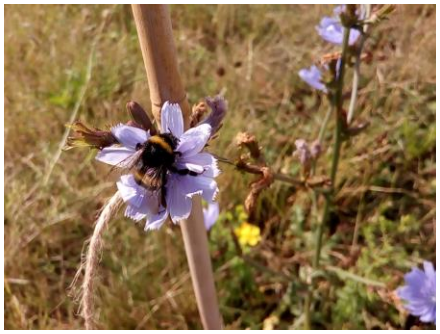</td><td>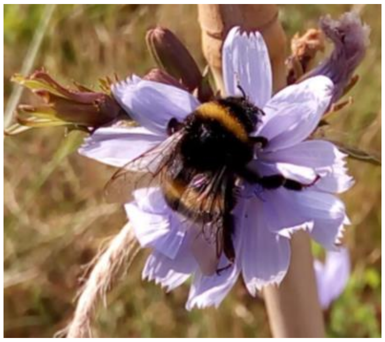</td></tr>
<tr><td colspan="2">Left: good enough capture; Right: this image is superior as it minimises background noise and provides an excellent top-down view of the insect. It might require being very close to the target flower though.</td></tr>
<tr><td>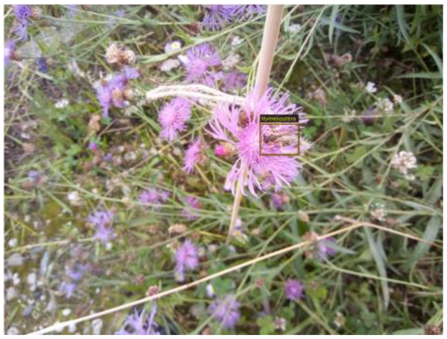</td><td>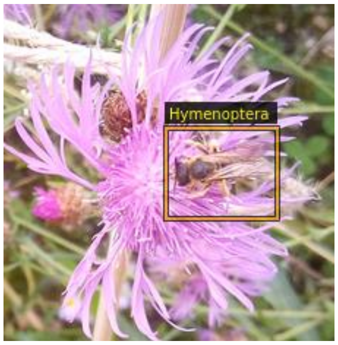
</td></tr>
<tr><td colspan="2">If we are too distant from the target flower, there will be too much background "noise". Many insects will end up out of focus, and they are on the background flowers. This can make insect identification challenging or impossible (left). Therefore, position the phone closer to the target flower (right).</td></tr>
<tr><td>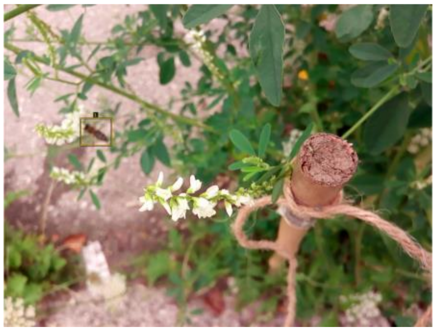</td><td>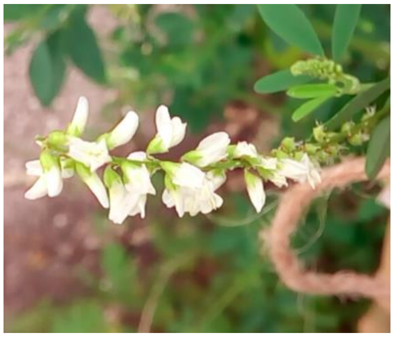</td></tr>
<tr><td colspan="2">The focus here is on the stick, not the main target flower (left). The stick should be beneath the flower and out of the camera's view (right). If you can't position the stick beneath the flower, avoid including the stick in the photo. Only the main flower should be visible with no other flowers in the background (right).</td></tr>
</table>

| | |
|---|---|
| 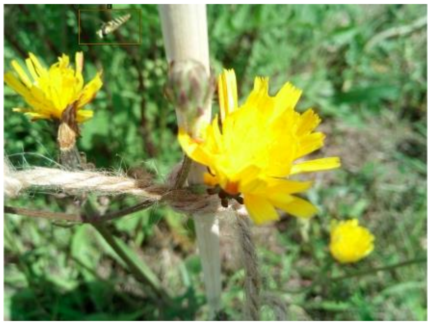 | 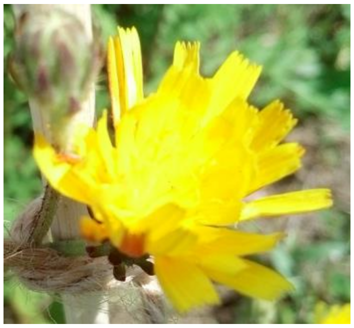 |
| Here, the focus is on the yarn, not the main flower. Be sure to dedicate enough time to lock the focus on the main flower. Avoid including the stick in the photo and avoid capturing background flowers that are out of focus. | |
| 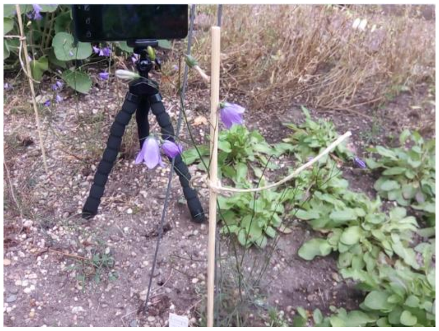 | 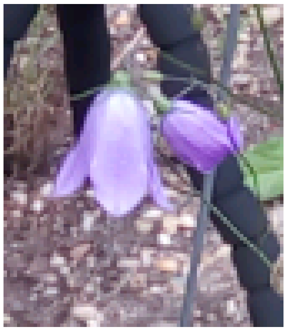 |
| Inflorescences are always a challenge. In this case, try to frame just one flower of the inflorescence, or a bunch, but not the entire inflorescence. As far as possible, avoid having artificial elements, like other phones, in the background. If you frame just one flower of the inflorescence, the insects will also occupy a larger area of the image. | |

#### B.5. Using yarn to secure the flower to the stick

Secure the flower to the stick with yarn without tying it too tightly. If the flower is bound too tightly with the yarn, it may stop the flow of fluids and tend to close, especially on sunny days.

The yarn should not be visible to the camera (see the image below for reasoning). If needed, use scissors to cut the yarn, or simply move the trailing yarn behind the flower(s), out of the camera's field of view.

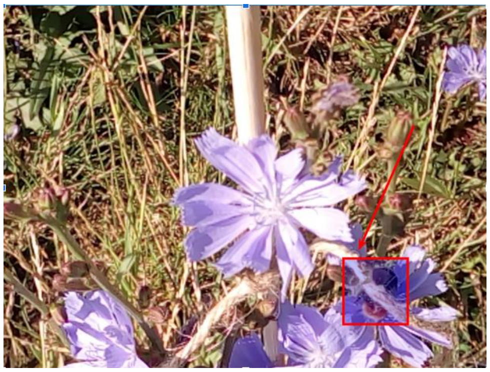

The bumblebee is obscured by the yarn - avoid such situations. The yarn should not be visible to the camera.

### C. Open Camera app guide

Link to the Android app - Open camera
(https://play.google.com/store/apps/details?id=net.sourceforge.opencamera&hl=en&gl=US )

You may also refer to the app's help page:
https://opencamera.sourceforge.io/help.html
https://opencamera.sourceforge.io/help.html#quickstart

#### C.1. Settings

Upon opening the app, locate a button resembling three vertically aligned dots at the top of the main display 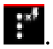. Clicking on this button will bring up a pop-up menu presenting the current settings (as shown in the screenshots below). "No Flash" and "Locked Focus" options should already be activated (their icons are underlined in the images below). The Photo Mode should be set to STD (Standard). Depending on your phone model, you may see different image resolutions.

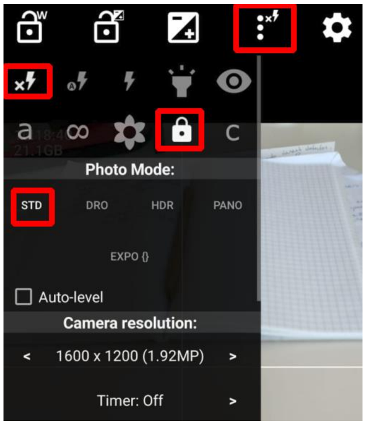

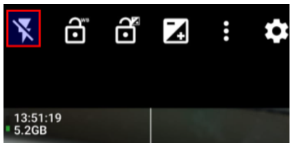

On some phones, you can check if the flash is off as shown above.

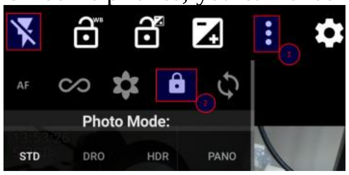

Clicking on the three vertical dots allows you to see if the lock icon is visible, indicating that the focus will remain locked during the time-lapse session.

To access all settings, use the Settings wheel/button 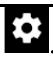. Here, the following settings should be enabled (as shown in the screenshot below):

- Face detection = off;
- Timer = off (no need for timer - this is for selfies);
- Repeat = unlimited (it takes photos until you click the shutter button again to stop);
- Repeat mode interval = 1s (attempts to take a photo every second).

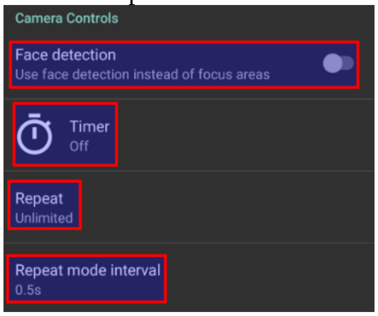

Under Settings wheel/button 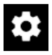 and 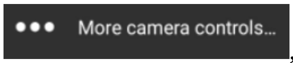

, set the following:

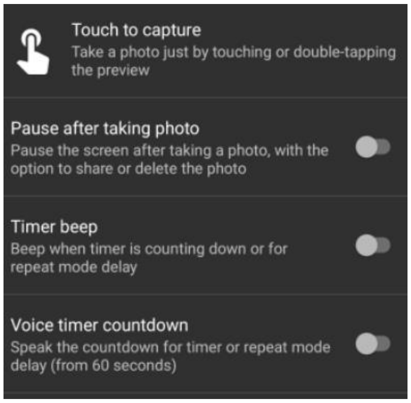

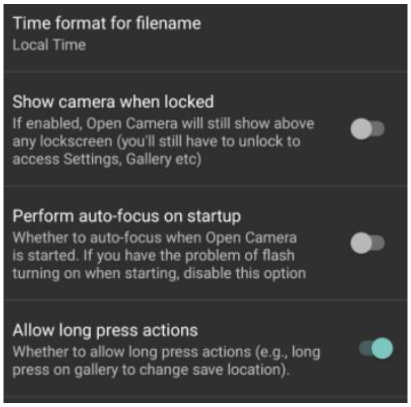

Under Settings wheel/button 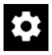 and 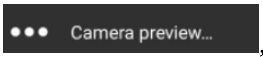

, set the following:

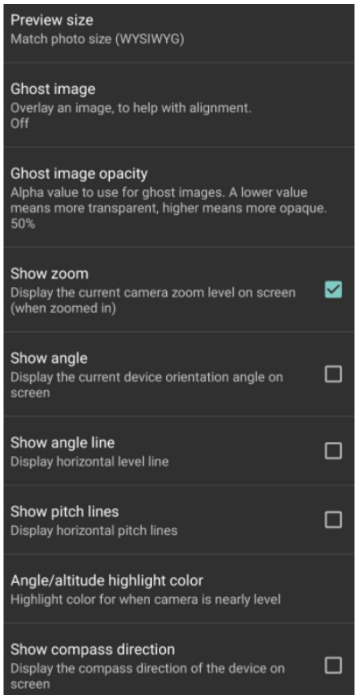

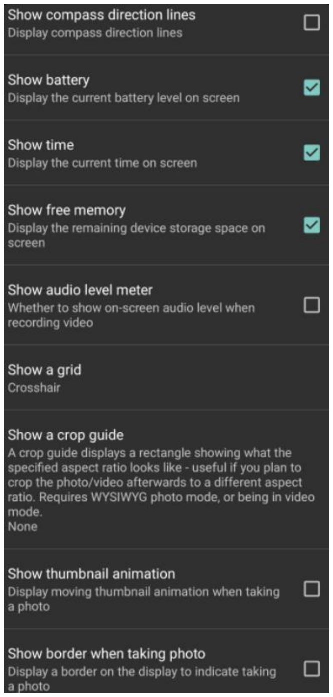

Under Settings wheel/button 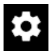 and 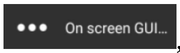

, set the following:

| | |
|---|---|
| 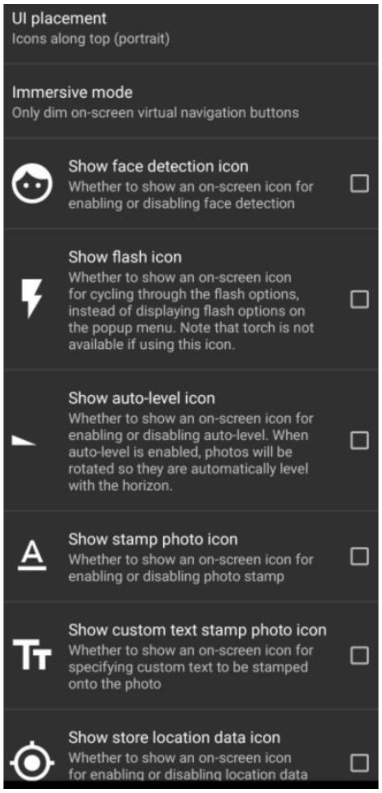 | 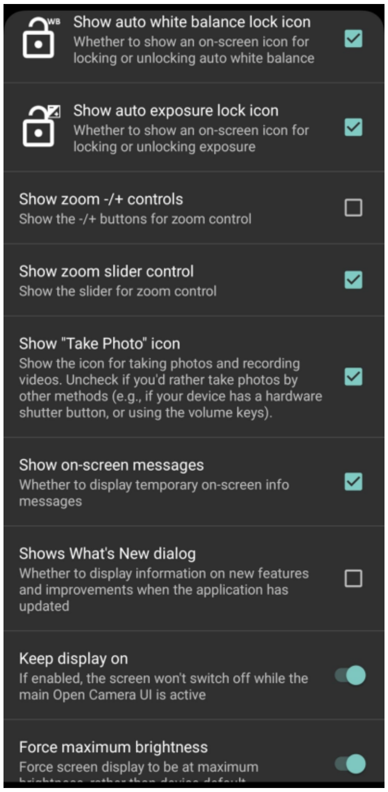 |

One crucial option to enable is "Keep display on". Despite its drain on battery life, we need to keep this on because Android otherwise forces the OpenCamera app into sleep mode. It's unclear if the developers will ever address this issue (see user forum discussion).

Under Settings wheel/button 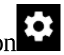 and 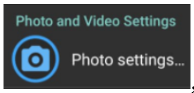

, set the following:

- Camera resolution - varies per phone model. Select option 1440 x 1080 (4:3, 1.56 MP); if it doesn't exist on your phone model, then opt for 1600 x 1200 (4:3, 1.92 MP).
  Note that even with the highest resolution, if the focus is blurred, resolution is not the limiting factor and we end up with many high-volume images that are slow for data transfer and provide no useful info. Thus, a good focus leading to sharp images of insects is far more important than image resolution.
- Image quality: 90% (default);
- Image format: JPEG (because it saves Exif metadata about ISO, aperture, phone, etc.; PNG doesn't and is converted from JPEG anyway);
- Save all images for HDR mode - OFF;
- HDR contrast enhancement - OFF;
- Exposure Bracketing Stops = 2 (default)
- Front camera mirror - OFF/No mirror (default);
- Copyright - leave empty
- Stamp photos - No stamp;
- Datestamp format = ISO 8601 yyyy-mm-dd;
- Time stamp format 24 h;
- GPS stamp format: Degrees/minutes/seconds
- Use addresses: Don't display address
- Distance unit: Meters
- Custom text: leave empty
- Font size: do not change (default 12)
- Font color: do not change (default white)
- Test style: do not change("Shadowed text" default)

C.2. How to create a “plant-folder”

For each flower, we take photos for 1 hour (you'll need to time this with a timer on your personal phone). For each session, you must create a different folder as follows: Settings wheel/button 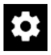 > More camera controls… > Save location > type the name of the plant, your initials (2 letters), and an ID number (for example: centaurea-scabiosa-vs-01), then press OK.

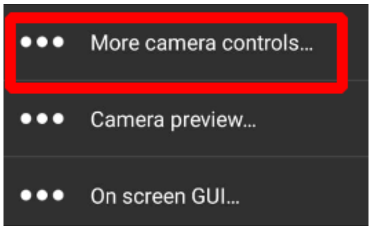

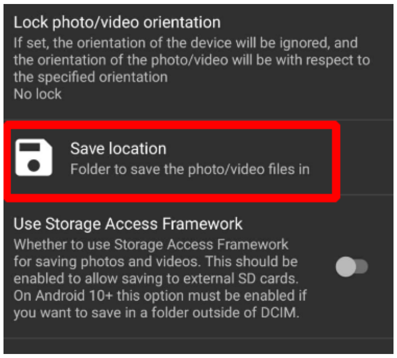

Each plant should have its own directory. Even if you have photographed a plant for one hour and then continue with the same plant species for the next hour, you must create a separate folder.

The name of the plant folder should include:

- The name of the plant species (you can use the Flora Incognita app to identify the plant). If you can only identify the genus, then you can use something like "centaurea-sp", or "fabaceae-sp" (using the plant family). If you can't identify it scientifically, then try to use two descriptive words, like "spiky-bushy". In any case, try to use two words separated by a hyphen.
- Your initials.
- An index: 01, 02, 03…, 09, 10, 11 and so on. For consistency, please use a zero before the digits smaller than 10.

The final plant-directory name should look something like this: "centaurea-scabiosa-vs-01", "centaurea-scabiosa-vs-02", and so on.
Please use hyphens (like this **-** ) instead of space as a separator. This is crucial for path management and data processing later on.
If it happens that you have the same initials as someone else, then communicate among yourselves so that each of you has a unique set of initials and uses them consistently.
Try to stick with just two letters in your unique initials setting (this ensures consistency in the file naming).

### D. Checklist after fieldwork day

#### D.1. Image download and upload

The overall workflow involves downloading the plant-folders from the phones to your laptop, and then uploading them to a backed-up storage space.

This data transfer process consists of two main steps. First, you download the plant-folders to your laptop. After that, you can upload them from your laptop to a secure storage. It's essential to do this in two steps because we've experienced data loss and interruptions when we tried to transfer data directly from the phones (phones can discharge or an accidental movement of the USB cable can cause the entire data transfer to fail and then you have to repeat it). You might generate 20-40Gb of images each day per smartphone.

##### D.1.1. Download plant-folders from the phones to your laptop

Firstly, download images from the phones directly to your laptop (you can use copy & paste to a location of your choice on your laptop). Keep in mind that this can take a considerable amount of time.

Downloading images from your phone to your laptop can vary depending on each phone type.

#### D.1.1.1. Connecting BlackView A60 smartphone models

When you connect their USB cables, a pop-up window will appear on the phone's main screen asking you to allow access for data transfer.

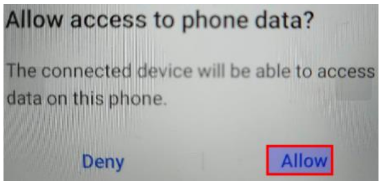

1. Connect the USB cable compatible with the phone to your laptop.

2. Locate the Settings wheel on the phone 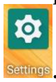. For the BlackView A60 smartphones, the settings wheel is also on the main screen. Go to "Connected devices", then "USB File Transfer" and then choose "File Transfer". This action will signal your laptop that the phone is ready for file transfer. Keep in mind that each smartphone model differs in its settings options.

3.

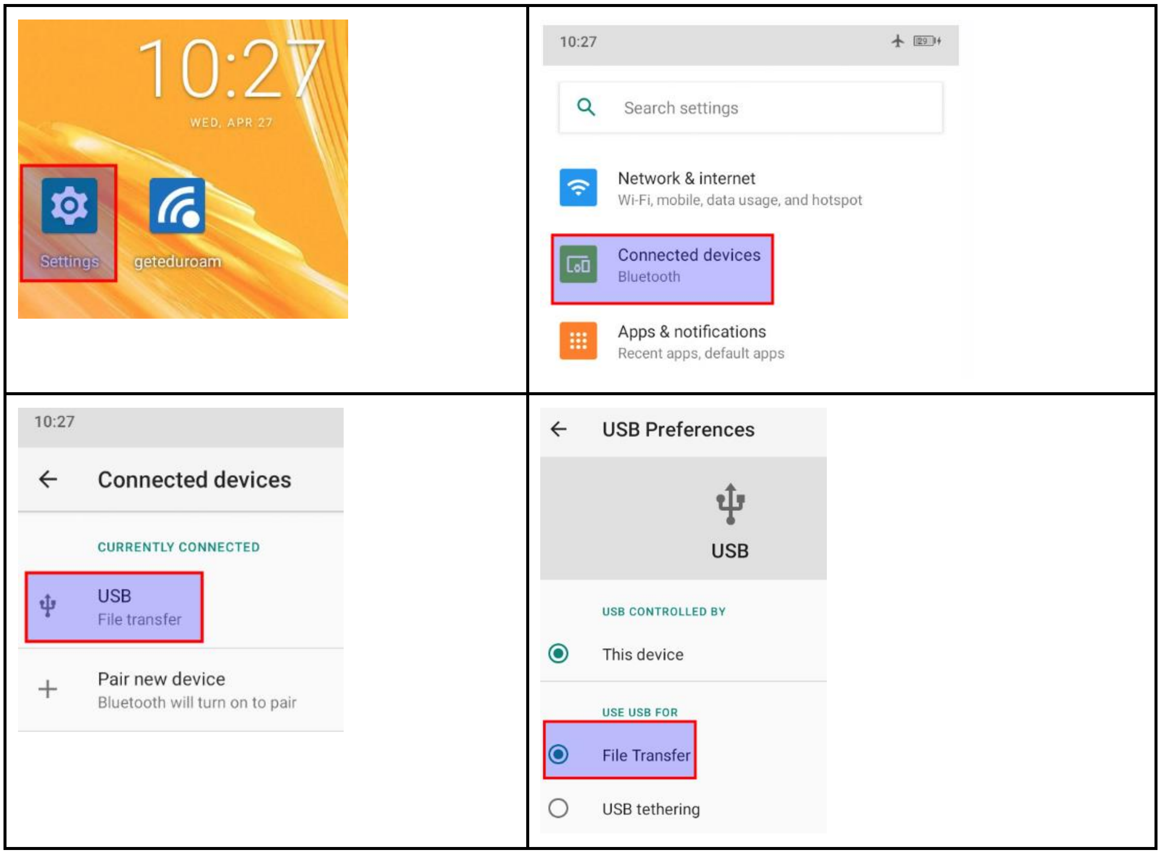

#### D.1.2.2. On Windows Operating System (OS)

Once your phone and laptop are connected (steps explained above), Windows will notify you that the phone is visible and ready for file transfer. You will see the phone name in the File Explorer in Windows (the File Explorer icon looks like a yellow folder and is usually located on the taskbar - lower-left corner of your screen). Navigate to the phone location, then to the DCIM directory where you should see the plant-folders that you created. Steps are illustrated below:

<table>
<tr><td>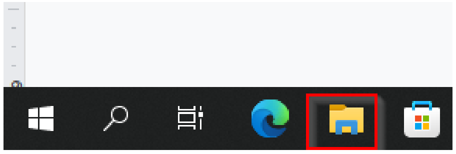</td><td>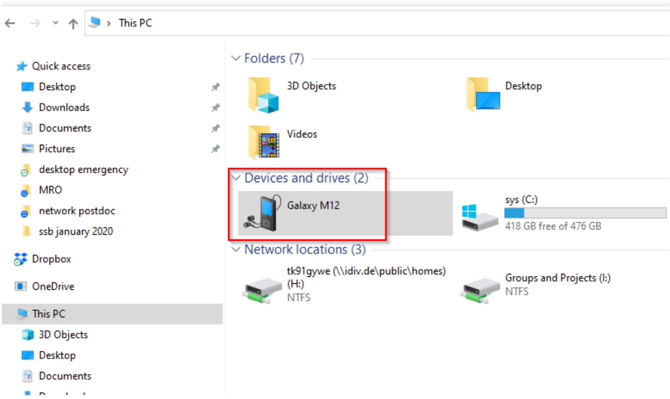
</td></tr>
<tr><td>1) File Explorer (for Windows). Click on it to visualise the content of your laptop.</td><td>2) Then, you should be able to see the name of your phone (names differ depending on the phone model). Double click on the icon corresponding to your phone to visualise the content.</td></tr>
<tr><td>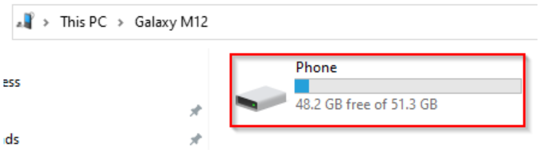
</td><td>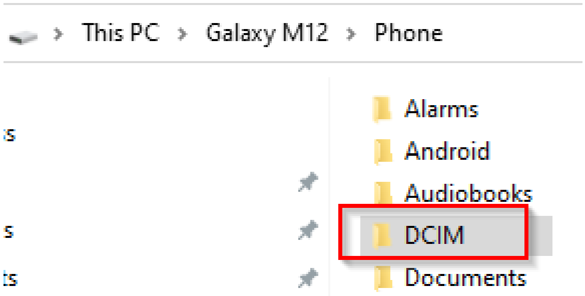
</td></tr>
<tr><td>3) Double click on the new icon.</td><td>4) Double click on the DCIM directory. Here Android stores images by default.</td></tr>
<tr><td>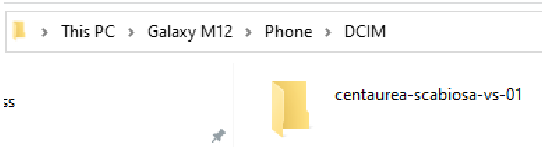
</td><td></td></tr>
<tr><td>5) Then you can see your plant folders. Each should contain thousands of images.</td><td></td></tr>
</table>

Now, create a directory on your laptop, in a location of your choice (e.g., in the "Documents" directory - navigate with File Explorer) with the exact date of the fieldwork day (we can call this the “date-directory” or “date-folder” in this tutorial). Please use the date format "year-month-day", like yyyy-mm-dd (e.g., 2022-04-28). See illustrated steps below:

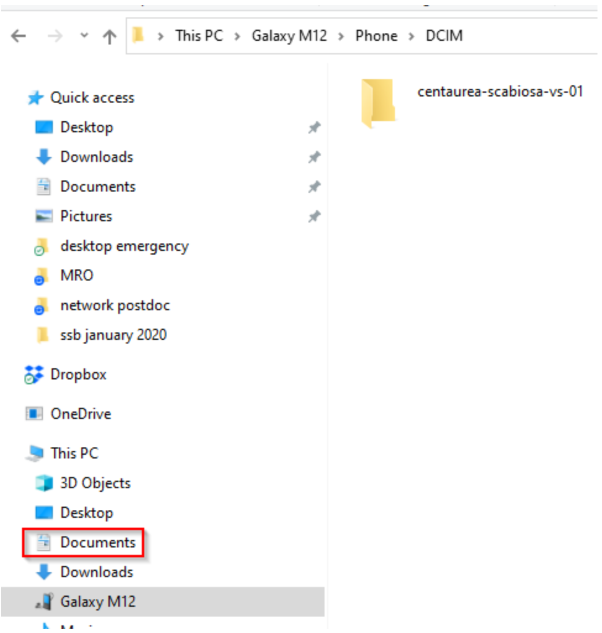

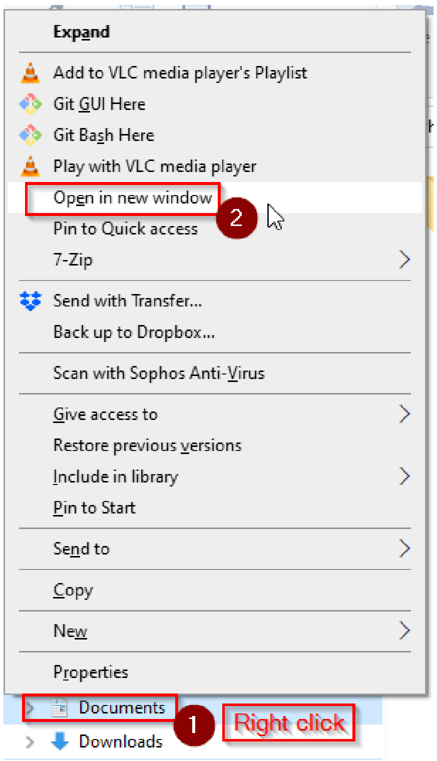

1) You find "Documents" on the left content list of the File Explorer. Right-click on "Documents" and a pop-up menu will appear, then select "Open in new window".

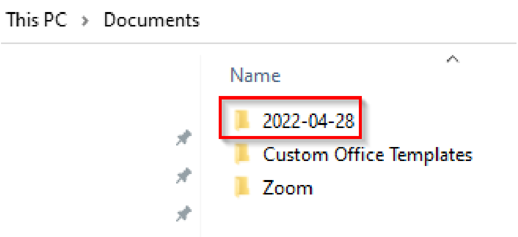

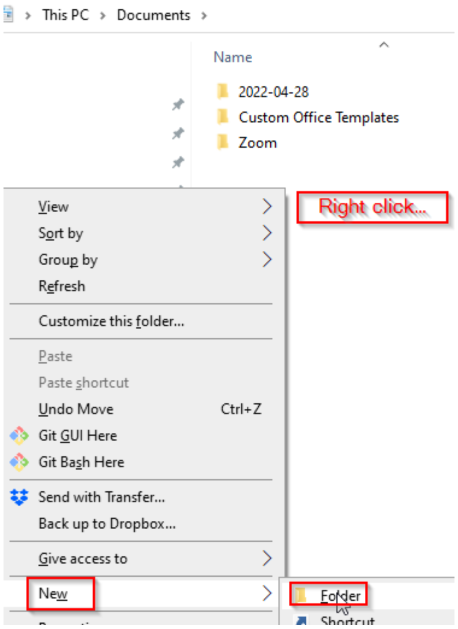

2) Create a date directory for that particular day. Respect the format yyyy-mm-dd.

To create a directory/folder: right-click on the empty content of the File explorer canvas, then choose New > Folder.

You can now copy the plant-folders from the phone to your laptop in the date-directory you created above.

Avoid having scattered images that do not belong to a plant-folder. Also, make sure to adhere to the details about the naming protocol in section C.2. "How to create a plant-folder". You should have done that directly in the field on the phones. This time, you can correct any typos while adhering to the naming convention.

Once the file transfer is completed (for each phone), connect the phones to charge along with the power banks connected to the electric grid (see section "D.4. Charge phones & power banks"). While they charge, you can now delete the plant-folders from the phone (see section "D.2. Delete the plant-folders from the phone").

Now is a good time to review your downloaded photos on your laptop and check if they align with the guidelines and examples given in sections "B.2. Selection of open flowers" & "B.4. Image background and distance from the flower". This provides feedback and you can learn what to improve for the next fieldwork day.

#### D.2. Delete the plant-folders from the phone

Each phone model varies, but the general procedure typically includes the following steps:

- Navigate to the Gallery / DCIM folder using the File Manager app.
- In the DCIM folder, you should see the plant-folders that you created during the day.
- Long press on a folder to activate the selection mode.
- Select the folder icons that you want to delete by clicking on each one.
- The delete option should appear.

Below is a step-by-step example:

| | |
|---|---|
| 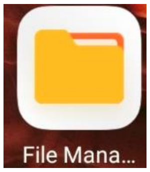  | 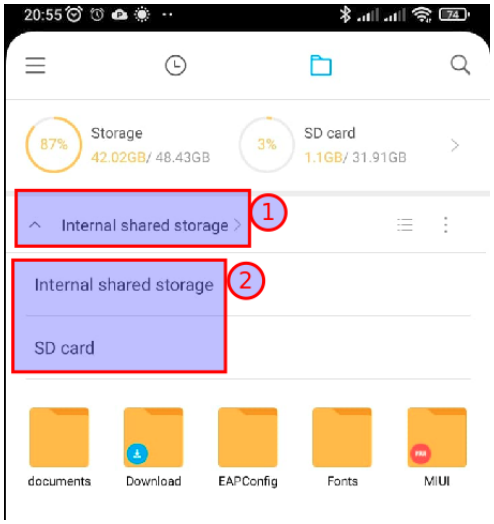  |
| 1) Locate the File Manager app on the phone (this varies from model to model). | 2) In this case, the OpenCamera app saves the plant-folders in the “Internal shared storage”. However, on some models, you might need to choose the SD card option. If you press on the arrow-up symbol, you should see two options (as shown in the image above). |

| | |
|---|---|
| 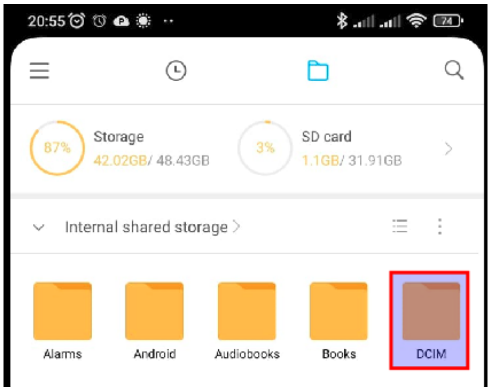 | 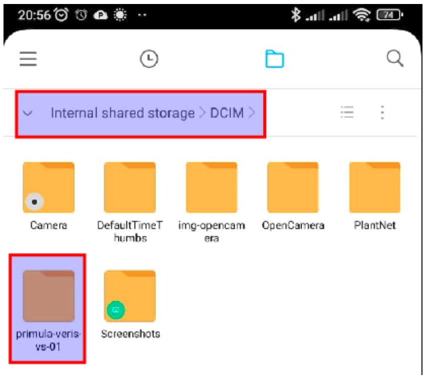 |
| 3) Navigate to the DCIM folder (this is where the Android OS usually stores photos). | 4) Inside the DCIM folder, you should see the plant-folders that you created with the OpenCamera app. If they are not there, check the OpenCamera folder (if it exists on your phone). Otherwise, the files might be stored on the SD card, so return to step 2 and choose the SD card path option. |
| 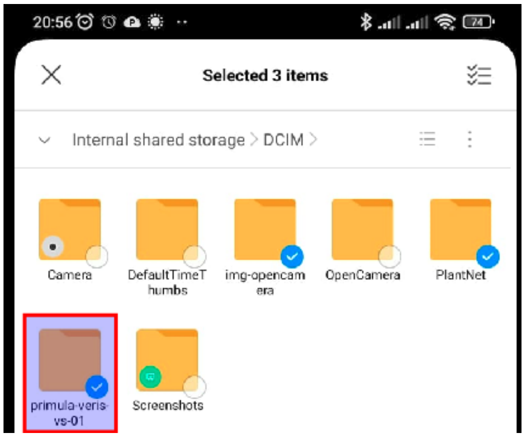 | 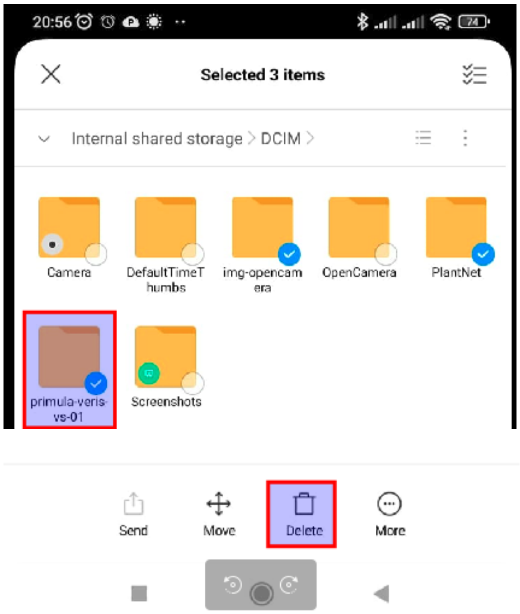 |
| 5) If you long-press on a plant-folder, Android will activate the multiple-selection mode (you'll see a transparent circle at the right corner of each folder). You can now lift your finger and touch each plant-folder that you want to select and then delete. Selected folders will have a blue circle with a "checked" symbol in the right corner. | 6) When you have selected all the plant-folders that you need to delete, the "delete" button should appear at the bottom of the screen. Press it, and the folders will be deleted. Be patient as this operation may take time and is likely irreversible due to its large size. |

#### D.3. Field notes

Here's an example of a site-info table that should be printed and used as field notes for each plant-folder, which means each time you set a phone to take photos for 1 hour:

| |
|---|
| Date (yyyy-mm-dd): |
| Site location (name/description): |
| GPS coordinates from Google maps/phone, e.g. 51.318204,12.396281 (latitude N, longitude E): |
| Observer name: |
| Plant-folder name (as given in your phone) |
| Device (see label on the back): |
| Weather (sunny/cloudy/rainy): |
| Time start recording (hh-mm): |
| Time end of recording (hh-mm): |
| I observed insects on the flower (yes/unsure) |
| Observations/Notes/Comments: |

At the end of the day or later, you need to digitise these field notes by filling in a corresponding Excel spreadsheet.

Each plant-folder (1 hour of observations) should have its own Excel file named "site-info" and contains the table from above. Name the spreadsheet to match its corresponding plant-folder name (e.g., centaurea-scabiosa-vs-01.xlsx). To rename a file, right-click on its icon and choose the rename option. Save the Excel file with its corresponding plant-folder under the corresponding date-folder.

#### D.4. Charge phones & power banks

It's recommended to turn off the phones whilst they charge to preserve battery life.

There may not always be enough chargers available to charge both the phones and power banks simultaneously. If this situation arises, first plug the power banks into the power source. Then, connect the phones to the charging power banks. The phones will slowly charge from the power banks, which in turn will be charging directly from the mains electricity (see image below):

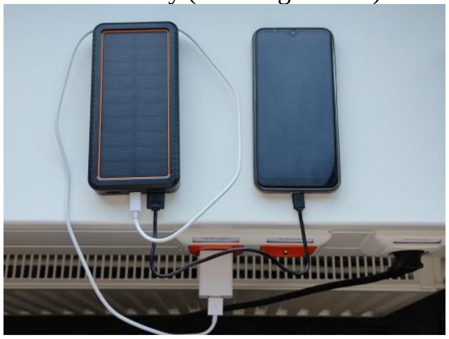

**Appendix Table VIII. List of sampling sites in and around Leipzig and Halle, Germany.**

Site metadata provided by Amibeth Thompson.

| Site name | Type | Latitude (WGS84) | Longitude (WGS84) | Notes |
|---|---|---|---|---|
| HAL-721-1 | Meadow | 51.489865 | 11.933866 | in park, gravel area, 125 m b/w transects |
| HAL-722-2 | Meadow | 51.490957 | 11.934477 | along road |
| HAL-723-3 | Meadow | 51.490471 | 11.939207 | in park |
| HAL-724-4 | Meadow | 51.489829 | 11.938118 | in park |
| HAL-725-5 | Meadow | 51.494844 | 11.949442 | in Peisnitz |
| HAL-726-6 | Meadow | 51.494526 | 11.948066 | in Peisnitz |
| HAL-727-7 | Meadow | 51.496376 | 11.939412 | in gravel area, 30 m b/w transects |
| HAL-728-8 | Meadow | 51.496914 | 11.938965 | in gravel area |
| HAL-729-9 | Meadow | 51.500061 | 11.942803 | between Wohnung, 75 m b/w transects |
| HAL-730-10 | Meadow | 51.500394 | 11.941824 | between Wohnung, 75 m b/w transects |
| LEI_A-701-1 | Roadside | 51.381921 | 12.272399 | along gravel spot |
| LEI_A-702-2 | Roadside | 51.381344 | 12.271475 | along road |
| LEI_A-703-3 | Meadow | 51.375605 | 12.279626 | meadow |
| LEI_A-704-4 | Meadow | 51.376405 | 12.278860 | meadow |
| LEI_A-705-5 | Meadow | 51.373104 | 12.285967 | meadow, 80 m b/w transects |
| LEI_A-706-6 | Meadow | 51.374005 | 12.285946 | along river/creek |
| LEI_A-707-7 | Forest | 51.367095 | 12.279839 | forest edge, 95 m b/w transects |
| LEI_A-708-8 | Forest | 51.367667 | 12.278746 | forest edge, 95 m b/w transects |
| LEI_A-709-9 | Meadow | 51.382265 | 12.266712 | along river |
| LEI_A-710-10 | Roadside | 51.381686 | 12.267349 | edge of meadow |
| LEI_C-837-1 | Roadside | 51.321546 | 12.385721 | NA |
| LEI_C-838-2 | Roadside | 51.321746 | 12.386616 | NA |
| LEI_C-839-3 | Meadow | 51.322031 | 12.389959 | NA |
| LEI_C-840-4 | Roadside | 51.326800 | 12.387749 | NA |
| LEI_C-841-5 | Roadside | 51.329163 | 12.382719 | NA |
| LEI_C-842-6 | Meadow | 51.320892 | 12.386599 | NA |
| LEI_C-843-7 | Meadow | 51.320296 | 12.397112 | NA |
| LEI_C-844-8 | Roadside | 51.321984 | 12.399551 | NA |
| LEI_C-845-9 | Roadside | 51.323747 | 12.405742 | NA |
| LEI_C-846-10 | Roadside | 51.323747 | 12.405742 | NA |
